# Oryza CLIMtools: A Genome-Environment Association Resource Reveals Adaptive Roles for Heterotrimeric G Proteins in the Regulation of Rice Agronomic Traits

**DOI:** 10.1101/2023.05.10.540241

**Authors:** Ángel Ferrero-Serrano, David Chakravorty, Kobie J. Kirven, Sarah M. Assmann

**Affiliations:** Biology Department, Pennsylvania State University, 208 Mueller Laboratory, University Park, PA, 16802, USA; Intercollege Graduate Degree Program in Bioinformatics and Genomics, Pennsylvania State University

**Keywords:** *Oryza sativa*, rice, G by E, adaptation, flowering time, GWAS, *OsHD2*, *OsSOC1*, heterotrimeric G protein, *OsRGA1*, *d1*, *OsGS3*, *OsDEP1*, *OsGGC2*

## Abstract

Modern crop varieties display a degree of mismatch between their current distributions and the suitability of the local climate for their productivity. To this end, we present Oryza CLIMtools (https://gramene.org/CLIMtools/oryza_v1.0/), the first resource for pan-genome prediction of climate-associated genetic variants in a crop species. Oryza CLIMtools consists of interactive web-based databases that allow the user to: i) explore the local environments of traditional rice varieties (landraces) in South-Eastern Asia, and; ii) investigate the environment by genome associations for 658 Indica and 283 Japonica rice landrace accessions collected from georeferenced local environments and included in the 3K Rice Genomes Project. We exemplify the value of these resources, identifying an interplay between flowering time and temperature in the local environment that is facilitated by adaptive natural variation in *OsHD2* and disrupted by a natural variant in *OsSOC1*. Prior QTL analysis has suggested the importance of heterotrimeric G proteins in the control of agronomic traits. Accordingly, we analyzed the climate associations of natural variants in the different heterotrimeric G protein subunits. We identified a coordinated role of G proteins in adaptation to the prevailing Potential Evapotranspiration gradient and their regulation of key agronomic traits including plant height and seed and panicle length. We conclude by highlighting the prospect of targeting heterotrimeric G proteins to produce crops that are climate resilient.

## Introduction

Rice is arguably the most critical food source on the planet as the dietary staple of over half of the world’s population (Bin Rahman and Zhang, 2023). The Green Revolution was crucial to promoting food security worldwide in the late twentieth century despite the concurrent increase in demand for food as a result of exponential growth of the world’s population (Dalrymple, 1986). As farmers began to prioritize high-yielding crops, they often abandoned local crop varieties (Frankel, 1974), resulting in the loss of genetic diversity over time that characterizes modern rice varieties (Khoury et al., 2022). Selective breeding practices in crops have primarily prioritized high-yielding but genetically uniform varieties, which can reduce adaptive potential (Østerberg et al., 2017). Therefore, as crop diversity decreases, so does their resilience to extreme weather events, such as heatwaves, droughts, and floods, which are becoming more frequent due to the already noticeable effects of climate change, and are predicted to decrease crop yields globally (Food and Agriculture Organization of the United Nations et al., 2018). To promote global food security, crop production can be stabilized by increasing genetic diversity (Renard and Tilman, 2019). A promising approach is to restore to modern elite varieties the lost genetic diversity present in local varieties (landraces) that are adapted to the local climate, and thereby foster improved yield stability in diverse conditions (McNally et al., 2009; Lasky et al., 2015; Faye et al., 2019).

The 3,000 (3K) Rice Genome Project has provided high-quality sequencing data for 3,010 rice genomes, obtained using Illumina-based Next-Generation sequencing (3,000 rice genomes project, 2014; Wang et al., 2018). Indica and Japonica are the two main subspecies of Asian cultivated rice and are the products of two separate domestication events from the ancestral species *Oryza rufipogon* (Kato, 1930; Sweeney and McCouch, 2007). The 3K dataset provides information on the genetic diversity of 658 Indica and 283 Japonica landraces grown in South-Eastern Asia (Figure 1), for which their collection sites are known and their population structure and history have been described (Gutaker et al., 2020). By studying local landraces, researchers can explore the factors that have shaped rice diversity and identify genetic variants and traits associated with the local environment that are suggestive of local adaptation. This information can aid the development of new high-yielding rice varieties better adapted to anthropogenically altered climates, to improve food security and sustainability.

**Figure 1.**
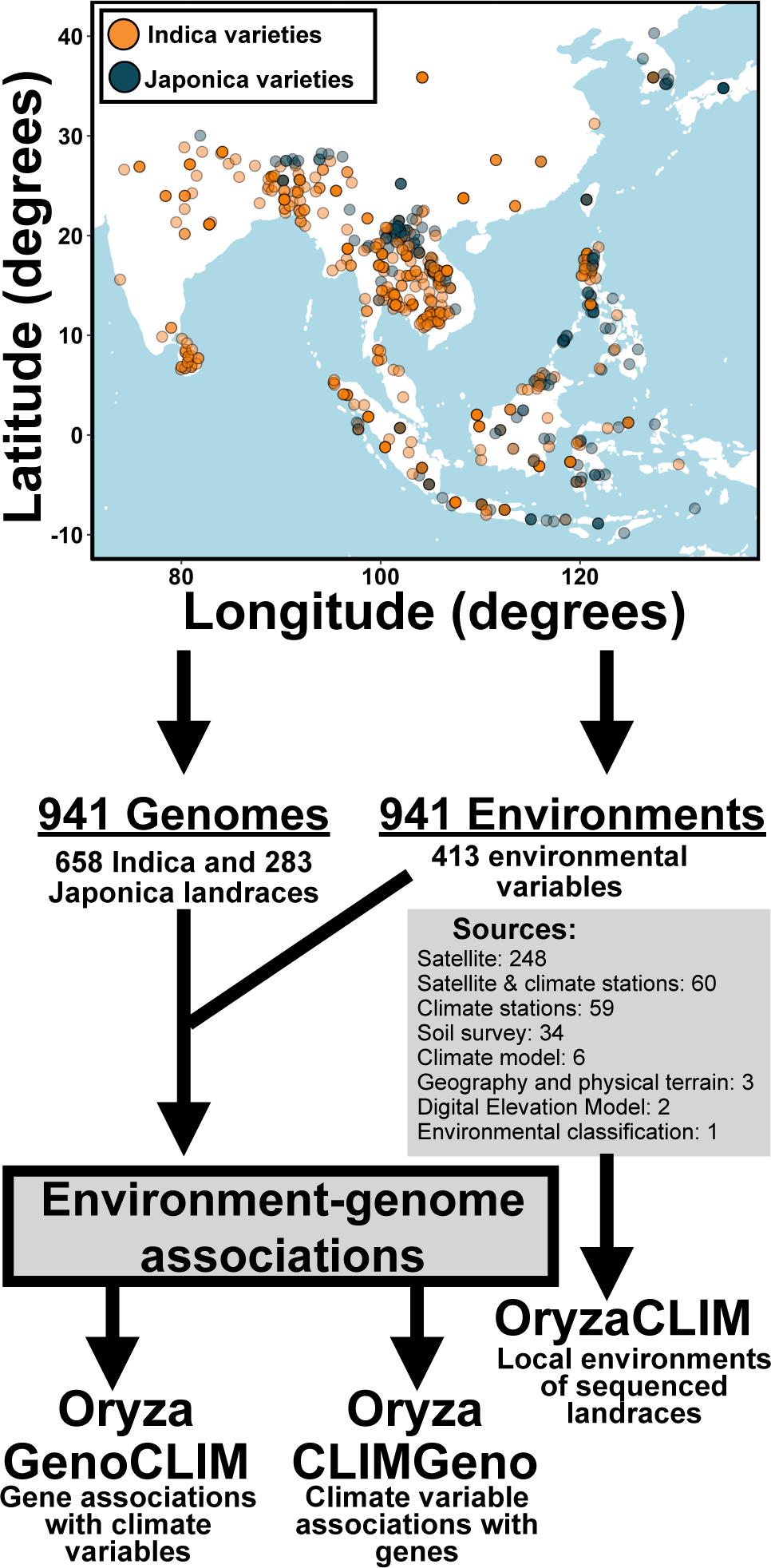
Global landrace distribution and environmental variables defining the local environment of the 941 Indica and Japonica fully-sequenced varieties used in this study to create the three components of Oryza CLIMtools. Geographic distribution of 658 Indica (orange data points) and 283 Japonica (blue data points) landrace varieties with known collection sites sequenced as part of the 3,000 (3K) Rice Genome Project (Wang et al., 2018; Gutaker et al., 2020). We curated 413 geo-environmental variables that describe the local environment of these landraces. Oryza CLIMtools provides a suite of online resources to provide gene by phenotype by environment associations to assist rice research and breeding programs.

Our recent work in locally-adapted accessions of *Arabidopsis thaliana,* collected from their local habitats, identified genome-wide associations between their natural genetic variation and a myriad of geo-environmental variables that define each local habitat. Arabidopsis CLIMtools provides a resource for the community to explore these associations (Ferrero-Serrano and Assmann, 2019; Ferrero-Serrano et al., 2022). However, similar resources have not been available for any crop species. Here, we present the first resource to provide gene by phenotype by environment associations to assist rice research and breeding programs. We integrate datasets comprising 413 geo-environmental variables that define the local environmental conditions at the collection sites of these 941 landraces and make this information available in OryzaCLIM (https://gramene.org/CLIMtools/oryza_v1.0/OryzaCLIM/). Using this environmental dataset, we establish an adaptive relationship between flowering time and the minimum temperature of the coldest month in Indica and Japonica landraces. Additionally, we present an interactive dataset of genotype–environment associations for 658 Indica and 283 Japonica rice landraces, with detailed information on the genes and variants statistically associated with the 413 environmental variables. These results are publicly available in a database platform, Oryza CLIMtools, available in the Gramene portal (Gramene.org; Tello-Ruiz et al., 2021). Oryza CLIMtools allows users to identify candidate adaptive genetic variants related to a given environmental variable of interest as codified in Oryza CLIMGeno and, conversely, to identify in Oryza GenoCLIM environmental variables associated with variation at any genetic locus of interest. We demonstrate the use of these tools, revealing an interplay among plant height, panicle length, and seed length that is associated with the Potential Evapotranspiration (PET) in the local environment and depends on both natural variation in the heterotrimeric G protein Gα subunit gene, *OsRGA1*, and a SNP that introduces a premature stop codon in the Gγ gene, *OsGS3*.

## Results

### Oryza CLIMtools

We created a comprehensive database of environmental data on rice landraces to serve as a valuable resource for researchers aiming to enhance rice research by integrating environmental parameters. This suite of online resources facilitates the process of incorporating environmental data into studies, and provides a user-friendly interface for researchers to access and utilize this information. Oryza CLIMtools (Figure 1) comprises three interactive web-based databases, OryzaCLIM, Oryza GenoCLIM, and Oryza CLIMGeno, that newly enable users to explore associations of promoter and transcript natural variants in 941 rice landraces, collected from their native range with 413 geo-environmental variables that define the environmental conditions in the areas where the landraces were collected (Figure 1). These environmental variables are collated in our OryzaCLIM tool and characterize the local environment of 658 Indica and 283 Japonica georeferenced and fully sequenced landrace varieties included in the 3K Rice Genome Project (Gutaker et al., 2020; Figure 1; Tables S1 and S2).

With Oryza CLIMtools (Figure 1), users can search for significant associations (FDR < 0.01), based on any climate parameter of interest using Oryza CLIMGeno. Alternatively, users can easily search for gene-climate associations (FDR < 0.01) based on any specific rice gene of interest using Oryza GenoCLIM. Of particular note, we also provide riboSNitch predictions, which indicate the likelihood that a variant alters RNA structure, which is a mechanism for differential post-transcriptional regulation (Halvorsen et al., 2010). As shown by our analyses in Arabidopsis (Ferrero-Serrano et al., 2022), riboSNitches are a fruitful topic for investigation.

We encourage the users of Oryza CLIMtools when using this resource to read the “considerations and limitations” document that we make available in Oryza GenoCLIM and CLIMGeno to familiarize themselves with the particularities of this approach and the impact that aspects such as population structure or the geographic distribution of the sampled population have in the interpretation our results.

Oryza CLIMtools has potential to assist the scientific community in ways that include:

Informing on the choice of cultivars for experimental GWAS analysis to facilitate inclusion of varieties from contrasting environments.

- Assigning relationships between phenotypic variability and environmental variation in this set of sequenced landraces, which can inform future studies of the functional interplay between genes and environments.
- Identifying the association of a particular environmental variable with variation in particular genes which, among other uses, may expedite forward genetics research.
- Identifying the association of genetic variation in specific genes of interest with environmental gradients which, among other uses, may expedite reverse genetics research.

We then illustrate the utility of these tools by: i) exploring the genetic basis of the interplay between flowering time and temperature in the local environment, and; ii) identifying a coordinated role of G proteins in adaptation to the local climate by regulating key agronomic traits.

### Natural variation in *OsHD2* and *OsSOC1* influences the flowering time by temperature relationship

To support the use of landraces to uncover the genetic basis of local adaptation, we asked whether there was an association between the phenotypic characteristics of these varieties and the environmental characteristics of the areas where they are grown. Using the environmental dataset we provide in OryzaCLIM, we uncovered a positive relationship in Indica and Japonica landrace varieties between flowering time as documented in publicly available data (Mansueto et al., 2017) and the minimum temperature of the coldest month (Karger et al., 2017) typical of their local environment (Figure 2). This relationship between flowering time and temperature is particularly strong in Japonica landraces (Figure 2). To uncover the genetic basis of this adaptive trait, we used Oryza CLIMGeno and identified in these Japonica landraces a haplotype consisting of a set of one missense and six intronic covarying SNPs (Figure 2A) in *HEADING DATE 2* (*OsHD2*/*OsPRR37*/*Ghd7.1*/*DTH7*; *Os07g0695100*), and the minimum temperature of the coldest month (Figure 2B). Landraces that contain the version of the haplotype in lower frequency in the Japonica population (minor allele) are typically grown in colder locations and display early flowering. By contrast, late-flowering landraces with the haplotype version more frequent in the Japonica population (major allele) are typically from warmer areas (Wilcoxon test, *p* < 0.0001; Figure 2B).

**Figure 2.**
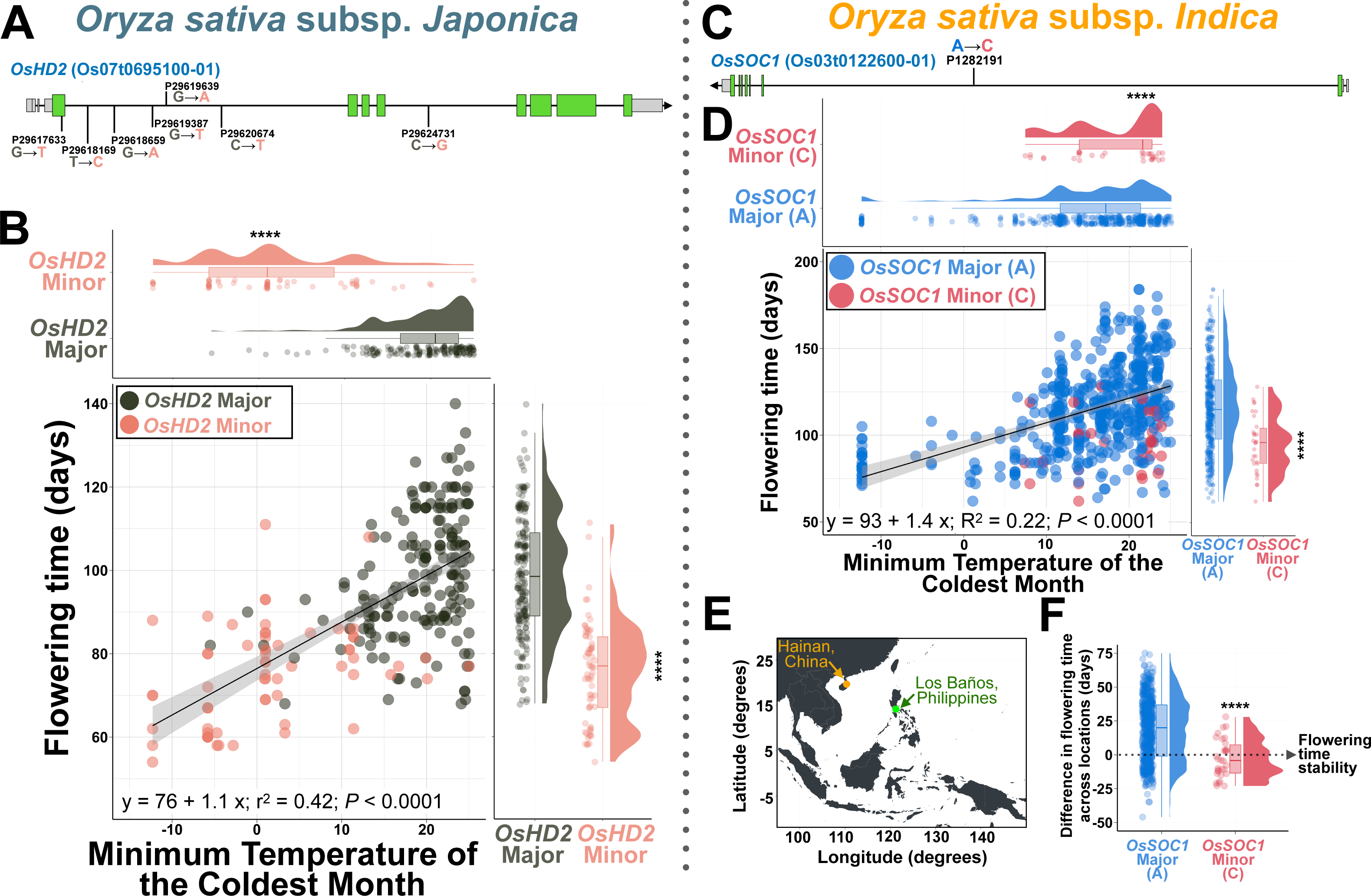
Natural variation in *OsHD2* tunes the relationship between heading date and temperature in the local environment; a minor variant of *SOC1* disrupts it. **A.** We identified in Japonica landraces an allele consisting of a set of one missense and six intronic covarying SNPs in *HEADING DATE 2* (*OsHD2*/*OsPRR37*/*Ghd7.1*/*DTH7*; *Os07g0695100*). Grey indicates UTRs, green indicates exons, and black indicates introns. **B.** Flowering time in Japonica landraces is significantly associated with the minimum temperature of the coldest month for their local environment (R^2^ = 0.42). Raincloud plots illustrate significantly different probability densities of the association of the local environment and flowering time for the major and minor alleles of *OsHD2* depicted in 2A. Coral datapoints and distributions represent landraces with the minor allele in *OsHD2*. Dark grey datapoints and distributions represent landraces with the major allele in *OsHD2*. **C.** We identified in Indica landraces an intronic variant in the floral pathway integrator and MADS domain transcription factor *SUPPRESSOR OF OVEREXPRESSION OF CONSTANS1* (*OsSOC1/AGL20/OsMAD50*; *Os03g0122600)* D. Flowering time in Indica landraces, similarly to Japonica landraces, is significantly associated with the minimum temperature of the coldest month for their local environment (R^2^ = 0.22). Raincloud plots illustrate significantly different probability densities of the association of the local environment and flowering time for the major and minor alleles of the intronic SNP in *OsSOC1* which is depicted in 3A. Red datapoints and distributions represent landraces with the minor allele in *OsSOC1*. Blue datapoints and distributions represent landraces with the major allele in *OsSOC1*. **E.** Planting sites in Hainan (China) and Los Baños (Philippines) for 576 Indica landraces with flowering time records at both of these different locations between these publicly available datasets retrieved from the Functional and Genomic Breeding (RFGB 2.0; Wang et al., 2020) and SNPseek (Mansueto et al., 2017) resource sites. **F.** Raincloud plot illustrating the significant flowering time stability of Indica landraces harboring the minor allele *OsSOC1* when planted at different locations. The difference in flowering time is calculated simply by subtracting the number of days it took landraces to flower in Hainan (China) from the days to flower in Los Baños (Philippines). The positive values for most landraces indicate that most landraces flowered later at lower latitudes in Los Baños, relative to those in Hainan. For the raincloud plots, in the box plots, the lower and upper boundaries indicate the 25th and 75th percentile, respectively. The whiskers below and above the box indicate the smallest value within 1.5 times the interquartile range (IQR) below the 25th percentile. The whisker above the box indicates the largest value within 1.5 times the IQR above the 75th percentile, respectively. The black line inside the box indicates the median. Pairwise nonparametric Wilcoxon tests were conducted to assess differences between alleles. **** (*p <*0.0001); *** (0.0001 < *p* < 0.001); ** (0.001 < *p* < 0.01); * (0.01 < *p* < 0.05); non-significant comparisons (ns) (*p >* 0.05). The associations between temperature and flowering time were fitted to a linear model. Regression lines are shown in black; gray shading represents 95% confidence intervals.

To uncover the genetic basis of flowering time adaptation to local temperature in Indica varieties, we used Oryza CLIMGeno and identified the association of an intronic variant in the floral pathway integrator and MADS domain transcription factor *SUPPRESSOR OF OVEREXPRESSION OF CONSTANS1* (*OsSOC1/AGL20/OsMAD50*; *Os03g0122600*; Figure 2C) with the minimum temperature of the coldest month in Indica landraces (Figure 2D). The minor allele of *OsSOC1* is found in landraces that are typically grown in warmer areas and exhibit early flowering (Wilcoxon test, *p* < 0.0001; Figure 2D). This relationship disrupts the overall strength of the association between flowering time and temperature in the local environment (R^2^ = 0.22, Figure 2B), which is stronger in the absence of the minor allele of *OsSOC1* (R^2^ = 0.26). From these findings, we hypothesize that the early flowering phenotype associated with the minor allele of *OsSOC1* may have been selected to favor flowering time stability and thus sustained yields across multiple locations. To test this hypothesis, we calculated the difference in flowering time for 576 Indica landraces with publicly available records resulting from two different planting locations (Figure 2E), one in Hainan (China) and the other in Los Baños (Philippines; Mansueto et al., 2017; Wang et al., 2020). The difference in flowering time between locations was calculated by subtracting the number of days it took landraces to flower in Hainan (China) from the days to flower in Los Baños (Philippines). In support of the conclusion that later flowering is typical of landraces collected from warmer climates (Figures 2B and 2D), we found that most landraces flowered later at the warmer location of Los Baños, relative to those in Hainan (Figure 2F). Additionally, we found that the flowering time of those landrace varieties harboring the minor *OsSOC1* variant remained remarkably stable when planted in these two different locations (Wilcoxon test, *p* < 0.0001; Figure 2F; note the reduced spread of datapoints around y = 0 for the minor *OsSOC1* variant, corresponding to flowering time homeostasis). We hypothesize that the minor OsSOC1 allele may have been selected by farmers to confer stability in flowering time, despite sacrificing adaptability to the local climate.

### Genetic variation in *OsRGA1*, the α subunit of the rice heterotrimeric G protein, is associated with a gradient in Potential Evapotranspiration

Potential Evapotranspiration (PET) is defined as a measure of the ability of the atmosphere to remove water through evapotranspiration given unlimited moisture (Thornthwaite, 1948; Hargreaves and Samani, 1982; Penman and Keen, 1997). Therefore, PET provides an estimate of the amount of water-use required for crop growth and can be used to help determine the appropriate amount of water that different varieties require to optimize productivity in a given location. We extracted the mean monthly PET of the coldest quarter (from now on, also referred to as “coldest quarter PET”), from the ENVIREM datasets (Title and Bemmels, 2018), for the local environments of the rice landraces utilized in this study. Using bioclimatic variables derived from monthly precipitation and temperature records (Karger et al., 2017) that we collated in OryzaCLIM, we determined that for both Japonica and Indica landraces, coldest quarter PET was co-associated with other temperature variables (Figure 3A). For example, areas with higher PET during the coldest quarter of the year are typically warmer areas with higher precipitation regimes (Figure 3A).

**Figure 3.**
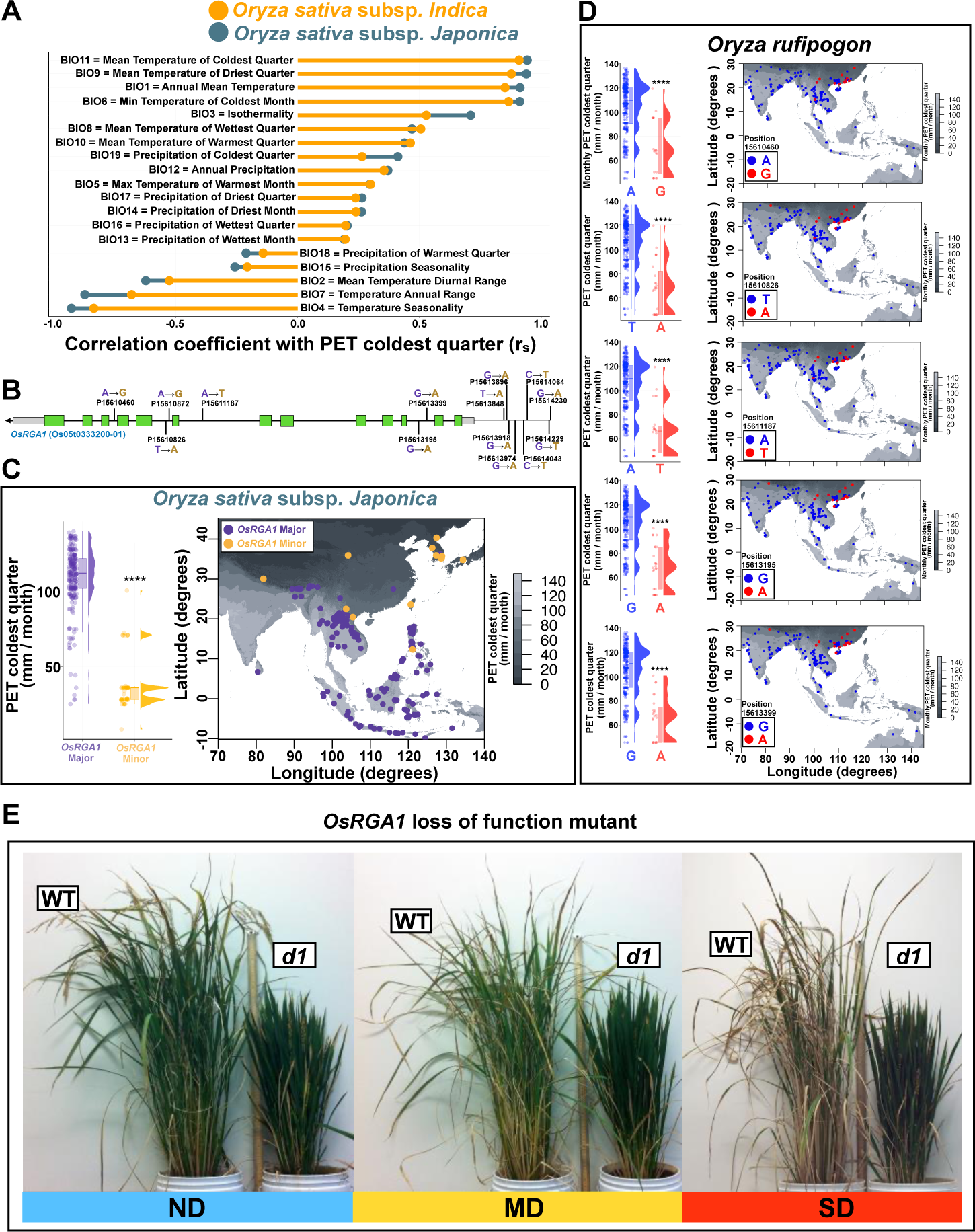
Haplotypic distribution of SNPs in *OsRGA1* is significantly associated with the mean monthly potential evapotranspiration (PET) during the coldest quarter of the year that is typical of their local environment. **A.** The mean monthly PET during the coldest quarter (“PET coldest quarter”) is co-associated with other climate variables. For example, areas with higher PET during the coldest quarter of the year are typically warmer areas with higher precipitation regimes **B.** We identified in Japonica landraces an allele consisting of six covarying intronic SNPs and eight upstream SNPs in *OsRGA1* (*Os05g0333200*) **C.** Major and minor alleles of *OsRGA1* are associated with the mean monthly PET during the coldest quarter. The map of South-Eastern Asia shows the geographical distribution of the major and minor haplotypic variants of *OsRGA1*. The grayscale color gradient in this map illustrates the cline in mean monthly PET during the coldest quarter. Purple dots and distributions represent landraces harboring the major allele in *OsRGA1*. Orange dots represent landraces harboring the minor allele in *OsRGA1*. **D.** Genetic variation present in 446 imputed sequences of *Oryza rufipogon* five of the six intronic SNPs uncovered in *OsRGA1* in Japonica landraces (positions 15610460, 15610826, 15611187, 15613195, 15613399) are also associated with mean monthly PET during the coldest quarter of the year in this wild rice species. Blue dots and distributions represent landraces with the major allele of *OsRGA1*. Red dots represent landraces with the minor allele in *OsRGA1*. The same colors depict the haplotypic distribution on the map of South-Eastern Asia with a color gradient that represents the cline in this drought-related variant. **E.** Assessment of drought tolerance of the *d1* mutant available in the Japonica cultivar Taichung 65 (T65) confirms its drought tolerance (see Figure S2). For panel A, r_s_ represents the Spearman’s rank correlation coefficient. For panels B and C: **** (*p* < 0.0001; non-parametric Wilcoxon test).

Heterotrimeric G proteins are signal transduction complexes that consist of Gα, Gβ, and Gγ subunits. In the classical model, upon activation of an associated G protein coupled receptor, the Gα subunit exchanges GDP for GTP, resulting in dissociation from the Gβγ dimer. Both Gα and Gβγ interact with intracellular effectors to evoke downstream signaling cascades. The system is returned to the inactive state when the intrinsic GTPase activity of Gα hydrolyzes GTP to GDP, thereby restoring the affinity of Gα for Gβγ and resulting in an inactive heterotrimeric complex (McCudden et al., 2005). The rice genome encodes one canonical Gα (*OsRGA1*), one Gβ (*OsRGB1*), and five Gγ (*OsRGG1*, *Os*RGG2, *Os*GS3, *Os*DEP1, and *Os*GGC2) subunits (Ueguchi-Tanaka et al., 2000; Utsunomiya et al., 2011). G protein subunits have been proposed to play a major role in regulating agronomic traits and stress responses (Botella, 2012; Cui et al., 2020). Our previous work has specifically implicated the rice Gα subunit, *OsRGA1*, in a number of agronomically-relevant traits, including plant architecture, drought sensitivity, light use-efficiency, and mesophyll conductance (Ferrero-Serrano and Assmann, 2016; Ferrero-Serrano et al., 2018; Zait et al., 2021).

We, therefore, used Oryza GenoCLIM to explore possible associations between previously uncharacterized natural variation in *OsRGA1* and the local environment of rice landraces. In Japonica landraces, we identified a correlation between a haplotype consisting of six covarying intronic SNPs and eight upstream SNPs in *OsRGA1* (*Os05g0333200*; Figure 3B), and PET during the coldest quarter (Wilcoxon test, *p* < 0.0001; Figure 3C). The allele in lower frequency is typically associated with locations with lower PET, while the allele in higher frequency is typically grown in areas with higher PET during the coldest quarter (Figure 3C).

To ascertain the ancestral origin of this G by association that we find in cultivated rice landraces in a wild relative, we identified the genetic variation present in 446 imputed sequences of *Oryza rufipogon* with known geographical origin using OryzaGenome 2.1 (Huang et al., 2012; Kajiya-Kanegae et al., 2021). We determined that five of the six intronic SNPs uncovered in *OsRGA1* associated with PET in Japonica landraces are also present in *Oryza rufipogon*, although not in strict covariation. We extracted the mean monthly PET of the coldest quarter from the ENVIREM datasets (Title and Bemmels, 2018), of these 446 wild rice lines and confirmed a similar pattern of association between natural genetic variation in *OrRGA1* and PET. Namely, *O. rufipogon* varieties that harbor these minor variants in *OrRGA1* are native to areas that experience lower PET. Conversely, *O. rufipogon* lines with the major variants experience higher PET (Wilcoxon test, *p* < 0.0001; Figure 3D). The fact that genetic variation in *RGA1* is associated with a gradient in coldest quarter PET across the local environment in a wild rice relative, and that not only the genetic variation, but also its association with the same environmental variable is conserved in rice landraces supports the hypothesis that these natural variants in *OsRGA1* are adaptive.

In the distribution range of the georeferenced landraces studied here, areas with lower PET typically experience lower temperatures and lower precipitation regimes. We, therefore, hypothesize that landraces grown in climates of lower PET during the coldest quarter (and lower precipitation; Figure 3A) should have been selected for drought tolerance. To evaluate this, we utilized the naturally occurring *d1* mutant available in the Taichung 65 (T65) Japonica cultivar, with a two bp deletion in *OsRGA1* that results in a protein null allele (Oki et al., 2009). *d1* and wild-type rice plants were kept well-watered for 60 days post-germination, after which soil was maintained at 100% (no drought, ND), 45% (moderate drought, MD), or 35% (severe drought, SD), relative water content (RWC). The dramatic drought-tolerant phenotype of *d1* is illustrated in Figure 3E. In the absence of drought (ND), both genotypes exhibited similar photosynthetic rates. However, under MD and SD, *d1* photosynthesis exceeds that of wild-type (Figure S2).

### A stop codon polymorphism in *OsGS3*, a rice G protein γ subunit, shows an association with a Potential Evapotranspiration gradient and interacts with *OsRGA1* to regulate vegetative and reproductive traits

Given our findings on *OsRGA1*, we next used Oryza GenoCLIM to explore the relationship between genetic variation in other G protein subunits and climate variables. We did not identify strong patterns of interaction with the environment in variants of the sole rice Gβ subunit gene, *OsRGB1 (Os03g0669200*). However, we uncovered a strong association between a polymorphism that introduces a premature stop codon to *OsGS3* (*Os03g0407400*), a Gγ subunit of the rice heterotrimeric G protein (Figure 4A), and the same gradient in mean potential evapotranspiration (PET) during the coldest quarter that we identified for *OsRGA1* (Figures 4B and 4C). Interestingly, within the sequenced Japonica population the full-length *OsGS3* allele (G) corresponds to the minor allele, while the allele containing the premature stop codon (T) represents the major allele (Figure 4A). We used OryzaGenome 2.1 (Kajiya-Kanegae et al., 2021) to explore the genetic variation of *GS3* present in 446 imputed sequences of *Oryza rufipogon* with known geographical origin (Huang et al., 2012). However, the position of *GS3* in *O. rufipogon* coincides with a region of low coverage for which no information is available, so we were unable to further investigate the presence of *GS3* SNPs in this wild-rice species.

**Figure 4.**
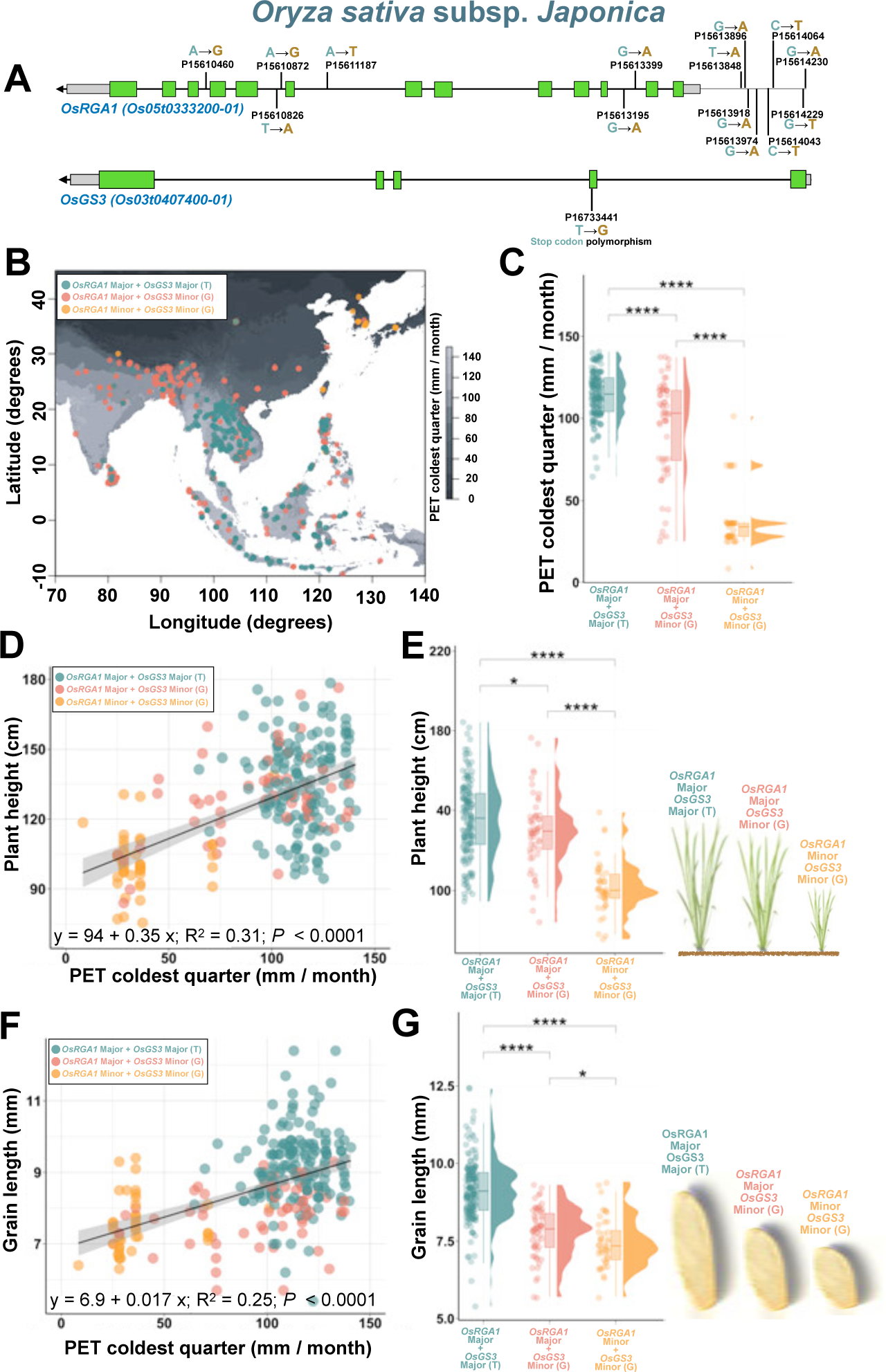
Haplotypic distribution of *OsRGA1* and a SNP introducing a stop codon in *OsGS3* in Japonica landraces are significantly associated with mean monthly potential evapotranspiration (PET) during the coldest quarter of the year and natural variation in plant height and seed length. **A.** We identified in Japonica landraces an allele consisting of six covarying intronic SNPs and eight upstream SNPs in *OsRGA1* (*Os05g0333200)* and a polymorphism (G to T) that introduces a premature stop codon in *OsGS3* (*Os03g0407400*). **B.** The map of South-Eastern Asia shows the geographic distribution of the three different *OsRGA1*-*OsGS3* allelic combinations that are present with allele frequency for both genes > 0.5 in Japonica landraces. The grayscale color gradient illustrates the cline in PET during the coldest quarter. **C.** Raincloud plots illustrate significantly different probability densities of the mean monthly PET during the coldest quarter for these three different *OsRGA1*-*OsGS3* allelic combinations. **D.** There is a significant association between the mean monthly PET during the coldest quarter and plant height. Allelic variation in *OsRGA1* determines this strength of this association and is strongly associated with plant height, as shown in Panel **E**, which is consistent with the well-reported dwarf phenotype of *OsRGA1* loss of function mutants (Oki et al., 2009; See Figure S2). **F.** There is also a significant association between the mean monthly PET during the coldest quarter of the year and grain length. Allelic variation in both *OsRGA1* and *OsGS3* determine the strength of this association, and minor variants of both genes are strongly associated with plants that display shorter grains, as shown in Panel **G**, consistent with the well-reported grain length phenotypes of *OsRGA1* and *OsGS3* mutants (Fan et al., 2006; See Figure S3). Pairwise nonparametric Wilcoxon tests were conducted to assess differences between alleles. **** (*p <* 0.0001).

We uncovered a significant association between landrace height in Japonica varieties and mean monthly PET during the coldest quarter of the year (R^2^ = 0.31; Figure 4D). We next found that shorter plants, which were typically found in areas with lower PET areas, were also more likely to simultaneously possess the minor *OsRGA1* allele and the minor (full-length) *OsGS3* allele (Wilcoxon test, *p* < 0.0001; Figure 4E), while taller plants from areas with higher evapotranspiration regimes typically harbor the major alleles of both *OsRGA1* and *OsGS3* (Wilcoxon test, *p* < 0.0001; Figure 4E). The *OsGS3* major allele, encoding a premature stop codon, has previously been named *GS3-3* (Mao et al., 2010); here we name it *OsGS3-3* for consistency. Landrace varieties from intermediate evapotranspiration regimes typically harbor the minor *OsGS3* allele in the major *OsRGA1* allele background (Wilcoxon test, *p* < 0.0001; Figure 4C). Although plant height in the landraces varieties with the minor *OsGS3* allele and the major *OsRGA1* allele was significantly shorter than those with the major *OsRGA1* allele and the *OsGS3-3* allele (Wilcoxon test, *p* < 0.05), the average height associated with the minor *OsRGA1* allele suggests that *OsRGA1*, not *OsGS3,* is the primary regulator of the variation in plant height in a G protein-mediated process across a PET gradient.

Using a publicly available dataset of agronomic traits that includes this landrace collection (RFGB 2.0; Wang et al., 2020), we also identified a significant association between landrace variety seed length and coldest quarter PET (R^2^ = 0.25; Figure 4F). Shorter grains are typical of landraces grown in low PET (drier and colder and less isothermal areas; Figure 4F), and these varieties are more likely to harbor simultaneously the minor *OsRGA1* allele and the minor *OsGS3* (full-length) allele (Figure 4G). Landraces with longer grains, grown in warmer, wetter, and less seasonal areas, typically harbor both the major allele of *OsRGA1* and the major *OsGS3-3* allele. Landrace varieties from areas with intermediate PET typically harbor the minor *OsGS3* allele in the major *OsRGA1* allele background. Although the average seed length in landraces with the minor *OsRGA1* allele and minor *OsGS3* allele is significantly shorter than in those with the major *OsRGA1* allele and the minor *OsGS3* allele (Wilcoxon test, *p* < 0.05), the magnitude of the average seed length associated with the introduction of the minor *OsGS3* allele implicates the Gγ (*OsGS3*), and not the Gα (*OsRGA1*) subunit as the primary regulator of the variation in seed length in this G protein mechanism across a PET gradient, notwithstanding an additional contribution by *OsRGA1* (Figure 4G).

### Genetic variation in the *OsGS3* and *OsDEP1* Gγ subunits in rice interact to determine seed and panicle length, but not plant height across a Potential Evapotranspiration gradient

In the continued evaluation of associations between natural variation in rice G proteins and PET, we highlight another Gγ subunit, in Indica varieties. A single non-synonymous variant in *OsDEP1* (Figure 5A) was found in association with the mean monthly potential evapotranspiration (PET) during the coldest quarter, the identical environmental parameter that we observed to be associated with *OsRGA1* and *OsGS3* variants in Japonica landraces. We found that the *OsDEP1* minor allele only occurred in combination with the *OsGS3* major allele (which in contrast to Japonica, is ‘G’, or the full-length *OsGS3* allele, in the Indica population), and landraces with this *OsDEP1* minor and *OsGS3* full-length combination are typically grown in areas with low PET (Wilcoxon test, *p* < 0.0001; Figures 5B and C). As expected, the long-grain *OsGS3-3* (non-full-length) allele (’T’) retained its association with the long-grain phenotype in Indica varieties (Wilcoxon test, p < 0.0001; Figure 5D). Variation in *OsDEP1* is only found in varieties with a full-length *OsGS3* allele and was not found in association with variation in grain length in the absence of variation in *OsGS3* (Wilcoxon test, *p* > 0.05; Figure 5D). This suggests that grain length is regulated by natural variation in *OsGS3* and not *OsDEP1*. Notably, the minor allele in *OsRGA1* (Figure 3B and 4A) was not present in a single one of the 658 Indica landraces, so there was no genetic interaction possible between *OsRGA1* and either *OsGS3* or *OsDEP1* within this Indica population.

**Figure 5.**
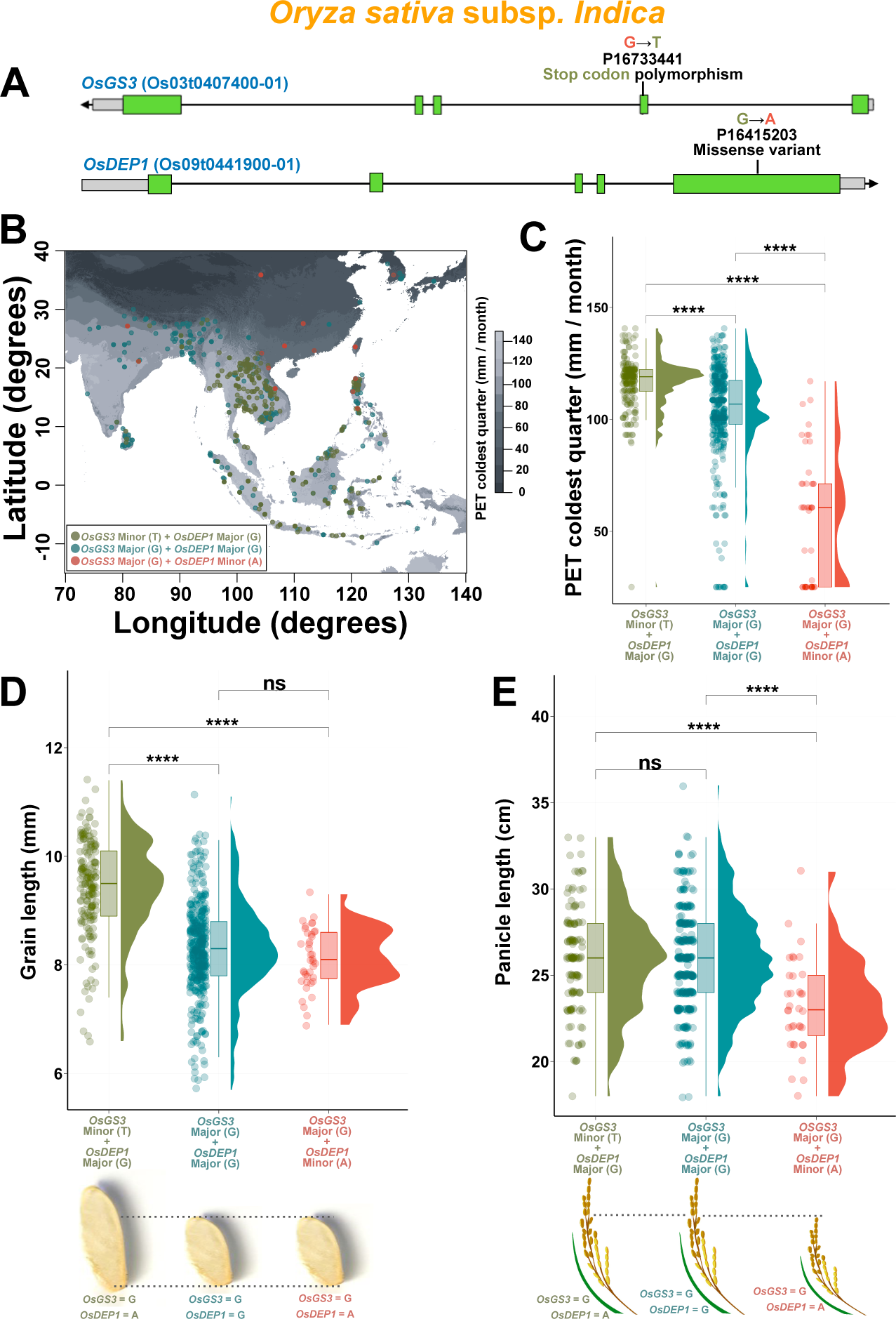
Haplotypic distribution of *OsDEP1* and *OsGS3* variants in Indica landraces are significantly associated with the mean monthly potential evapotranspiration (PET) during the coldest quarter of the year and natural variation in seed and panicle length. **A.** We identified in Indica landraces a stop codon-causing (G to T) SNP in *OsGS3* and a missense variant in *OsDEP1*. **B.** The map of South-Eastern Asia shows the geographic distribution of the major and minor variants of the different *OsGS3*-*OsDEP1* allelic combinations. The grayscale color gradient illustrates the gradient in mean monthly PET during the coldest quarter (“PET coldest quarter”). **C.** Raincloud plots illustrating the association of the major and minor alleles of *OsGS3* in the presence/absence of the minor allele *of OsDEP1* with PET coldest quarter. Both B and C illustrate the association of the full-length (major; G) *OsGS3* allele and the minor *OsDEP1* allele with lower PET regions. **D.** Raincloud plot illustrates the association of *OsGS3* major (G) vs. minor (T) variants with grain length and the lack of association of the *OsDEP1* major vs. minor variants with grain length. **E.** Raincloud plot illustrates the association of the *OsDEP1* major vs. minor variants with panicle length and the lack of association of *OsGS3* major vs. minor variants with panicle length. Pairwise nonparametric Wilcoxon tests were conducted to assess differences between alleles. **** (*p <* 0.0001).

When we evaluated the association of *OsGS3* and *OsDEP1* with panicle length, obtained from a publicly available dataset that includes this landrace collection (RFGB 2.0; Wang et al., 2020), we found that Indica varieties with the minor *OsDEP1* allele had shorter panicles relative to the major allele, regardless of the presence of *OsGS3* variation in this major *OsDEP1* allele background (Wilcoxon test, *p* < 0.0001; Figure 5E). This suggests that panicle length is regulated by natural variation in *OsDEP1* and not *OsGS3*.

To support the observation that variants in the Gα subunit, *OsRGA1*, and not the Gγ subunits regulate plant height, as we observed in *OsRGA1* and *OsGS3* Japonica lines (Figure 4E), we explored the existing variation in plant height across the different *OsGS3*- *OsDEP1* allelic combinations in these Indica landraces. As all Indica landraces harbor the major *OsRGA1* allele, the lack of association between plant height and any combination of *OsGS3-OsDEP1* alleles (Wilcoxon test, *p* < 0.05; Figure S3), confirms that natural variation within these Gγ subunits does not influence plant height in Indica landraces.

### Agronomic traits regulated by natural variation in G proteins that affects the nature of their protein-protein interactions

As described above, we used Oryza CLIMtools to uncover associations of G protein natural variants in Indica and Japonica landraces with the local environment and with adaptive agronomic traits. We hypothesize that our observation that natural variation in *OsDEP1* and not *OsGS3* regulates panicle length in Indica, while *OsRGA1* and *OsGS3* do so in Japonica (Wilcoxon test, *p* < 0.0001; Figure S4) may be explained by: i) the absence of the *OsRGA1* minor allele in Indica, and; ii) the different versions of a third Gγ subunit, *OsGCC2*, that we observe in Indica and Japonica. In *OsGGC2*, a set of three variants in the last exon plus one in the 3’ UTR covary, defining one allele that is categorically found in all Japonica landraces in CLIMtools, and another allele that is categorically found in Indica landraces in CLIMtools (Figure 6A). Given that rice Gγ subunits are known to function both additively and antagonistically (Sun et al., 2018), a resultant difference in *Os*GGC2 function may shift the equilibrium of signaling from other Gβγ dimers, and thereby alter the genetic interactions between subunits.

**Figure 6.**
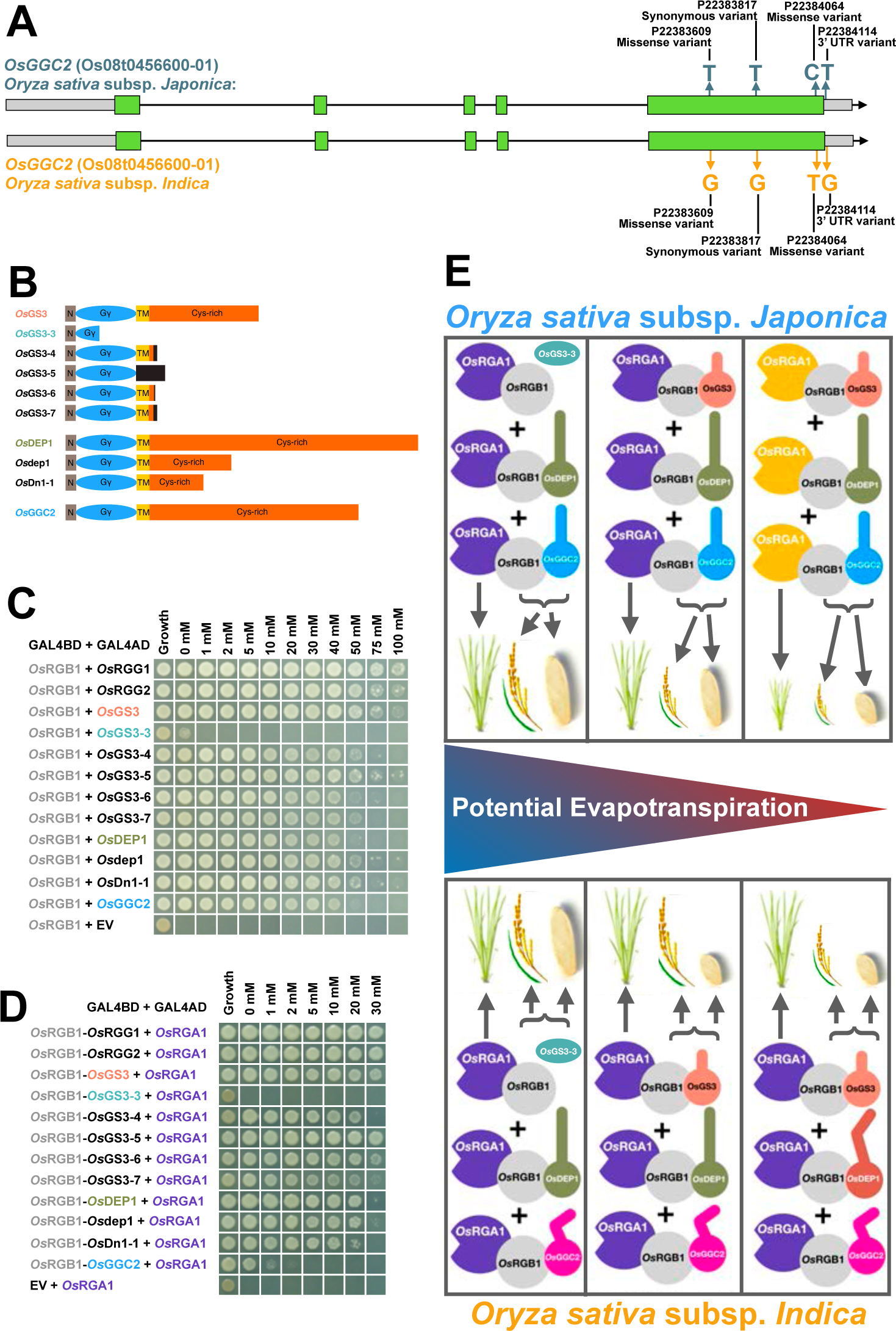
Genetic variation in heterotrimeric G proteins is associated with a gradient in the mean monthly potential evapotranspiration (PET) during the coldest quarter of the year, and the interplay of vegetative and reproductive traits. **A.** We uncover a set of three covarying variants in the last exon plus one in the 3’UTR of *OsGGC2* with one allele typical of Japonica, and the other typical of Indica varieties, that may affect the equilibrium of signaling by other Gβγ dimers. **B.** Domain schematic of rice Type C Gγ subunits indicating known truncations. N = N-terminal sequence, Gγ = Gγ-like domain, TM = putative transmembrane domain, Cys-rich = C-terminal Cys-rich domain; the dark gray boxes correspond to out-of frame translated regions. See Figure S5A and S5B for multiple alignments of protein sequences. **C.** Yeast 2-hybrid assays between Gβ (*Os*RGB1) and Gγ show lack of interaction between *Os*RGB1 and *Os*GS3-3 and positive interaction with all other Gβγ combinations. **D.** Yeast 3-hybrid assays between Gβγ dimers and the canonical Gα, *Os*RGA1, show that all *Os*RGG1, *Os*RGG2, *Os*GS3, *Os*DEP1 and *Os*GGC2 proteins encoded by Gγ alleles were able to complex with *Os*RGB1 and *Os*RGA1 via *Os*RGB1, except *Os*GS3-3. Assays in **C.** and **D.** were spotted on SC-Trp-Leu media as a growth control, and SC-Trp-Leu-Met-His with the concentrations of 3-Amino-1,2,4-triazole (3-AT) indicated above each column to assess interaction strength. 3-AT is a competitive inhibitor of the *HIS3* reporter gene of the yeast 2-/3-hybrid systems and therefore allows for semi-quantitative assessment of interaction strength, with growth at higher 3-AT concentrations indicating a stronger interaction. **E.** Cartoon summarizing our hypothesis on how the alternative combinations of the different heterotrimeric G protein variants associated with mean monthly PET during the coldest quarter of the year affect agronomic traits. Color-coding for Gγ alleles is as in panels B-D. Color-coding: *Os*RGA1 purple = major allele and yellow = minor allele, *Os*RGB1 = gray, *Os*GS3 salmon = full length allele and turquoise = *Os*GS3-3 truncated allele, *Os*DEP1 olive green = major allele and orange-red = minor allele, *Os*GGC2 light blue = Japonica allele and pink = Indica allele. See Discussion for details.

We have previously shown that a C-terminal truncation of the Arabidopsis ortholog of *Os*GS3, *At*AGG3, that retains only part of the Gγ domain (residues 1-78), loses interaction with the Arabidopsis Gβ subunit (Chakravorty et al., 2011). Therefore, we hypothesized that the similarly truncated protein encoded by the Japonica major allele/Indica minor allele (T) of *OsGS3*, *OsGS3-3* (Figure 6B), would also not bind the rice Gβ subunit. Using the yeast 2-hybrid assay of protein-protein interaction, we observe that the protein encoded by the *OsGS3-3* allele indeed displays a severely impaired interaction with *Os*RGB1, as evident by weak yeast growth that occurs only in the absence of 3-aminotriazole (3-AT; Figure 6C). By contrast, all full-length Gγ subunits, corresponding to the Japonica Taichung 65 sequences of *Os*RGG1, *Os*RGG2, *Os*GS3, *Os*DEP1, and *Os*GGC2, strongly bound *Os*RGB1 as assessed by growth on a semi-quantitative 3-AT concentration curve (Figure 6C). The previously identified truncations resulting from indels in *OsGS3* or *OsDEP1* that retain a full Gγ domain (Figure 6B), namely *OsGS3-4* (Mao et al., 2010), *OsGS3-5*, *OsGS3-6*, *OsGS3-7* (Takano-Kai et al., 2013), *Osdep1* (Huang et al., 2009) and *OsDn1-1* (Taguchi-Shiobara et al., 2011; alignments in Figure S5A and S5B), also all bound *Os*RGB1 with similar or increased strengths compared to the corresponding wild-type subunits (Figure 6C). As expected, no Gγ subunits displayed any growth on interaction selective media in combination with the empty vector control (negative control; Figure S5C). We also utilized a yeast 3-hybrid approach to assess the interaction of the different Gβγ dimers with Gα subunits. Consistent with our yeast 2-hybrid data, the truncated *Os*GS3-3 protein did not allow heterotrimer formation with *Os*RGA1 (Figure 6D). The full-length Taichung 65 Gγ subunits all allowed heterotrimer formation with *Os*RGA1 (Figure 6D). The variant Gγ proteins *OsGS3-4*, *OsGS3-5*, *OsGS3-6*, *OsGS3-7*, *Osdep1*, and *OsDn1-1* also allowed heterotrimer formation (Figure 6D). These results indicate that the *Os*GS3-3 allele likely acts as a functional null due to its inability to complex with other G protein subunits. Contrastingly, the Gγ truncations that can bind Gβ may act as dominant negatives, or sequester Gα and Gβ subunits since we have previously shown that the Cys-rich tail of the Arabidopsis ortholog of *Os*GS3, *At*AGG3, is required for Gγ function (Chakravorty et al., 2011). We did not examine the interactions of the proteins encoded by the *OsDEP1* minor allele (*Os*DEP1^C261Y^) or the *OsGGC2* Indica allele (*Os*GGC2^L169W^ ^R321C^) with *Os*RGB1, or with *Os*RGA1 when contained within the Gβγ dimer: the changes encoded within these Gγ protein variants occur in the Cys-rich tail, a region of Type C Gγ subunits that does not contribute to binding of other heterotrimeric G protein subunits (Chakravorty et al., 2011) and the tail is proposed to reside in the apoplast (Wolfenstetter et al., 2015). In Figure 6E, we synthesize all our G protein-related findings to propose a molecular mechanism by which rice G proteins’ natural variants regulate signaling, with phenotypic consequences (see Discussion).

### CLIMtools allows broad exploration of the genetic basis of local adaptation in rice

Above, we have demonstrated the utility of Oryza CLIMtools to uncover adaptive associations between phenotype and environmental variables codified in OryzaCLIM. We have used these tools to focus on agronomic roles of variation in heterotrimeric G proteins. It is unavoidable that our dataset may miss alleles that are only present in geographic locations where georeferenced landrace data are not available. However, as exemplified in Figure 7, Oryza CLIMtools can be utilized broadly to investigate many putative mechanisms for potential genetic bases of adaptation to diverse climate variables, including various temperature measures (Wilcoxon test, *p* < 0.0001; Figure 7 A-C, E,F), soil quality (Wilcoxon test, *p* < 0.0001; Figure 7D), illumination conditions (Wilcoxon test, *p* < 0.0001; Figure 7G), and aggregating variables such as latitude (Figure 7H).

**Figure 7.**
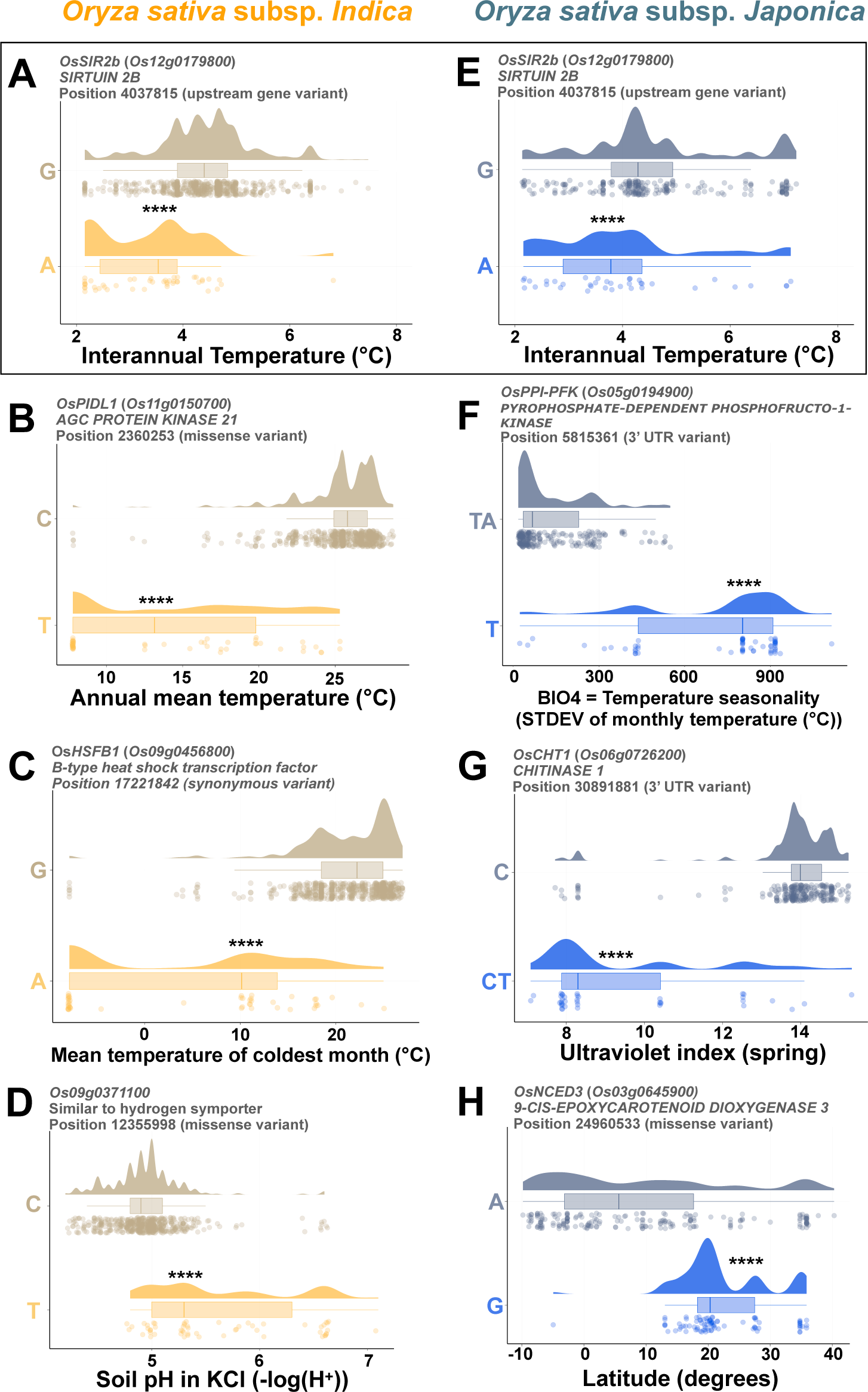
Examples of genotype by environment associations found using Oryza CLIMtools. Example associations for Indica landraces between **A**. an upstream gene variant in *OsSIR2b* and interannual temperature, **B.** a missense variant in *OsPIDL1* and annual mean temperature, **C.** a synonymous variant in *OsHSFB1* and the mean temperature of the coldest month, **D.** a missense variant in *Os09g0371100* and soil pH. In Japonica landraces, we uncovered an association between **E.** the same upstream variant that we identified in Indica landraces in *OsSIR2b* with the same climate parameter, interannual temperature, **F.** a 3’UTR INDEL variant in *OsPPI-PFK* and temperature seasonality, **G.** a 3’UTR INDEL variant in *OsCHT1,* and the ultraviolet index during the spring season, and **H.** a missense variant in *OsNCED3* and latitude.

## Discussion

As rice is the staple food for more than half of the world’s population and the main source of calorie intake in emerging and developing economies (Seck et al., 2012), any reduction in global production will have important implications for food security worldwide (Godfray et al., 2010). A major challenge is thus to feed a burgeoning human population while mitigating crop losses incurred as a result of climate change. There is already extensive information on the genetic diversity of rice, but the environment is the missing piece of the puzzle. To date, no comprehensive resource has been available to investigate the relationship between genetic and environmental variation that exists in the local environments where rice landraces are grown. With this in mind, we created OryzaCLIM (https://gramene.org/CLIMtools/oryza_v1.0/OryzaCLIM/), an intuitive online resource available to the community to explore 413 geo-environmental variables that define the local environmental conditions at the collection sites of 658 Indica and 283 Japonica sequenced landraces (Figure 1).

We then evaluated several significant (FDR < 0.01) climate-gene associations found using Oryza CLIMtools, to both illustrate the utility of our resources and exemplify how these associations can be validated to demonstrate causality.

### Flowering time and temperature

Average global temperatures are predicted to increase by 1.5°C between 2030 and 2052 (Delmotte et al., 2018). These rising temperatures pose a significant threat to rice production (Zhao et al., 2016) and flowering is, notably, the developmental stage most sensitive to heat stress (Matsui et al., 2001; Jagadish et al., 2010). Indeed, supraoptimal temperatures increase spikelet sterility and critically reduce rice yield (Shrestha et al., 2022).

Many species in natural settings demonstrate shifts in flowering time associated with ongoing climate change (Fitter and Fitter, 2002). In the case of rice, modern high-yielding cultivars are grown in many areas of the world that diverge enormously in their environmental conditions. The flowering time of these modern cultivars is not optimized for all local environments. By contrast, local landraces are cultivated varieties that have evolved as a result of both natural and artificial selection (Casañas et al., 2017) and are therefore amenable to mining for adaptations of flowering time to the local environment.

Temperature is a key environmental factor patterning contemporary rice genomic diversity (Gutaker et al., 2020). Indica and Japonica landrace varieties show considerable phenotypic variability in their flowering time (Figure S6A), and here we show that flowering time in both Indica and Japonica landraces is significantly associated with the minimum temperature of the coldest month in their local environment (Figures 2B and 2D). Rice is a facultative short-day plant. Many landraces in tropical and subtropical regions have mild photoperiod sensitivity that inhibits flowering under long days and promotes flowering under short days, maximizing the time for vegetative growth before flowering (Zong et al., 2021). At the same time, suboptimal temperatures negatively affect survival and yield (Zhang et al., 2014), and varieties from areas where temperatures during the cold season are suboptimal sacrifice vegetative growth but ensure early flowering and seed production before the onset of low temperatures concurrent with shorter photoperiods. In our study, we describe the relationship between flowering time and temperature in the local environment for a fully sequenced rice population. Introducing flowering time stability, as provided by the minor allele in *OsSOC1* (Figure 2D and 2E), may provide a farmer-desired trait that outweighs its climate incompatibility.

Our dataset covers a vast geographical range of 6,730 km in longitude and 3,388 km in latitude (Figure 1). However, it does not provide coverage that is comprehensive and geographically dense in areas such as China and other higher latitude regions, due to the lack of georeferenced landrace data from those regions. Previous studies that have focused on those regions have identified *OsHD1* and *OsGhd7* (Han et al., 2016; Zhang et al., 2015) as genes that are major regulators of flowering time in response to photoperiod. In our datasets, natural variants in these photoperiod-responsive genes in significant association with latitude can also be identified (*p* < 0.0001; Document S1), confirming previous work that focused on datasets from high latitude regions (Han et al., 2016; Zhang et al., 2015).

Besides photoperiod, temperature is a major regulator of flowering time transition and in this regard we identified an adaptive association between an allele in *OsHD2* and the minimum temperature of the coldest month (Figure 2B). *OsHD2* has been identified as a flowering time QTL that is important for temperature regulation of photoinduction (Nakagawa et al., 2005). The Arabidopsis ortholog of *OsHD2*, *AtPRR7*, is a key component of the temperature compensation mechanism to maintain a constant circadian rhythm under changing temperatures (Salomé et al., 2010). Consistent with our findings, previous work has shown that functional alleles of *OsHD2*, known to produce higher yields, are found mainly in warmer southern latitudes, while ‘weaker’ and early flowering, non-functional alleles of *OsHD2* are found in colder, northern latitudes (Yan et al., 2013; Koo et al., 2013; Li et al., 2015; Ye et al., 2018). These results are also consistent with the findings of Guo et al. (2020), who determined that varieties from areas with lower local temperatures are enriched in *OsHD2* allelic variants that are associated with temperature-sensitivity of flowering time. Conversely, varieties from areas with warmer temperatures harbor *OsHD2* alleles that are associated with temperature-invariant flowering times.

Since environmental variables often correlate among themselves (Figure S1), it is possible that other co-correlated variables also contribute to adaptation in flowering time to the local environment. Indeed, we observe that the minimum temperature of the coldest month correlates with other temperature and precipitation seasonality variables (Figure S7).

Given this finding, we propose *OsHD2* as a key mediator of the adaptation of flowering time to the local environment. The feasibility of fine-tuning flowering time via genome editing of the upstream open reading frame of *OsHD2* has also been demonstrated (Liu et al., 2021). We can envision an approach by which the appropriate flowering time of any given variety can be modulated to match local temperatures, to create varieties that are better adapted to their local climate and maximize their potential yield in a given location. Oryza CLIMtools opens the door to exploring the genetic basis of flowering time adaptation so that flowering time can be tailored to match environmental conditions in the area of cultivation.

### Genetic variation in heterotrimeric G protein Gα subunit *OsRGA1* is associated with a gradient in Potential Evapotranspiration

Most modern, high-yielding rice varieties rely on a mutant *SD-1* gene, introduced during the Green Revolution, as the basis of their semi-dwarf phenotype (Hargrove and Cabanilla, 1979; Hargrove et al., 1980; Ferrero-Serrano et al., 2019). However, the *qDTY1.1* allele, a QTL for drought tolerance, has been lost in these modern varieties due to its tight linkage in repulsion with the *sd1* allele. Consequently, the widespread use of *sd1* as a source of dwarfism both reduces the genetic diversity of this trait and results in modern rice cultivars that are typically drought sensitive. This illustrates one of the most documented consequences of genetic erosion in crops: the reduction in adaptability or tolerance to abiotic factors (Khoury et al., 2022).

Drought is one of the most widespread and damaging environmental stress factors (Farooq et al., 2012; Gupta et al., 2020) and affects rice particularly as varieties are often adapted either to rain-fed or fully irrigated systems. For this reason, the identification of an alternative source of dwarfism that also provides an adaptive advantage to drought can contribute to yield stability in global yield production. In this study, we argue that natural variants in other dwarfing genes are available that are positively associated with drought-related environments.

Following this rationale, we explored the association of natural-occurring variants in the heterotrimeric G protein Gα subunit, *OsRGA1,* and the local environment, particularly drought-related environmental variables. Naturally occurring functional null *OsRGA1* mutants display a characteristic dwarf phenotype (Ashikari et al., 1999; Oki et al., 2009; Ferrero-Serrano and Assmann, 2016; Zait et al., 2021). Loss-of-function *OsRGA1* dwarf mutant plants display improved drought resistance compared to wild-type plants as a consequence of their increased water use efficiency, increased mesophyll conductance, lower leaf temperature, and increased light use efficiency (Ferrero-Serrano and Assmann, 2016; Ferrero-Serrano et al., 2018; Zait et al., 2021).

Using Oryza GenoCLIM (https://gramene.org/CLIMtools/oryza_v1.0/Oryza_GenoCLIM/), we identified a set of covarying SNPs in the intronic and upstream regions of *OsRGA1* (Figure 3B) in association with a gradient in mean monthly Potential Evapotranspiration (PET) during the coldest quarter of the year. The minor allele is found in landraces collected from areas with lower PET, and the most frequent allele (major allele) is present in landraces collected from areas with higher PET (Figure 3C).

Potential evapotranspiration provides an estimate of the amount of water-use for growth and can therefore be used to select drought-tolerant varieties. As shown in Figure 3A, areas with lower PET during the coldest quarter correspond to colder but also more variable temperatures throughout the year and lower precipitation regimes. In other words, areas with lower PET are associated with more challenging and unpredictable environments (Figure 3A). We hypothesize that those landraces from areas of lower PET will have improved abiotic stress tolerance to multiple factors, a trait in which G proteins play a critical role (Wu and Urano, 2018; Wang and Botella, 2022). We illustrate this point by showcasing the improved drought tolerance of the *OsRGA1* natural null mutant, *d1* (Figure 3E).

The role of the minor *OsRGA1* variant in adaptation to regions with lower PET during the coldest quarter is supported by our finding that five of the six intronic *OsRGA1* variants that we found in rice landraces are conserved in the ancestral species of rice, its wild rice relative *Oryza rufipogon* (Huang et al., 2012), where they are also associated with a gradient in PET (Figure 3D). This finding in an ancestor of domesticated rice supports the hypothesis that natural variation in *OsRGA1* is adaptive across a PET gradient in South-Eastern Asia.

### Genetic variation in heterotrimeric G proteins is associated with a potential evapotranspiration (PET) gradient and the interplay of vegetative and reproductive traits

Previous studies typically describe the agriculturally-relevant roles of individual G protein subunits and their variants in isolation (Botella, 2012; Wu et al., 2022; Ferrero-Serrano and Chakravorty, 2023). Here, we reveal how interaction among natural variants in the Gαβγ complex determines agronomic traits and stress responses that contribute to fitness in the local environment. While we did not identify variants of interest in *OsRGB1*, we found different natural variants of the Gγ subunits in association with PET, the same environmental variable associated with the *OsRGA1* variants, suggesting a genetic interaction.

Gγ proteins are divided into three distinct groups according to their C-terminal domains, with the rice genome encoding two shorter Gγ subunit proteins, *Os*RGG1 (Type A) and *Os*RGG2 (Type B), and three longer Gγ subunits with a Cys-rich tail: *Os*DEP1, *Os*GGC2, and *Os*GS3 (Type C; Trusov et al., 2012). A series of indels has been identified in *OsGS3* and *OsDEP1* (Figure 6B) that confer changes in grain shape (Mao et al., 2010; Takano-Kai et al., 2013) and a dense and erect panicle morphology (Huang et al., 2009; Taguchi-Shiobara et al., 2011), respectively.

In Japonica rice, the minor allele of *OsRGA1* is associated with extremely short grains, significantly shorter than those conferred by the major *OsRGA1* allele in combination with the minor (full-length) *OsGS3* allele (Figure 4G). The role of *OsRGA1* may be via impairment of *OsDEP1* and *OsGCC2* signaling. Indeed, knocking out either *OsDEP1* or *OsGCC2* results in short plant, panicle, and grain phenotypes that are epistatic to *OsRGA1 d1* mutations (Sun et al., 2018; Chaya et al., 2022). In Indica rice, the *OsGS3* allele frequency is the reverse of that seen in Japonica (Figure 4B vs. 5B). In Indica landraces, the minor allele of *OsGS3*, previously named both *gs3* and *GS3-3*, is the allele that encodes a premature stop codon (T) within the Gγ domain. This allele was found to cause a long-grain phenotype (Fan et al., 2006; Mao et al., 2010) that we now show is found in areas of higher PET (Figures 4B and 4C). The *OsGS3-3* variant was introduced from a Japonica ancestor(s) into the Indica gene pool (Takano-Kai et al., 2009; Mao et al., 2010), which may account for the fact that the *OsGS3-3* allele is at a higher frequency in the Japonica population relative to the Indica population.

The *Os*GS3-3 protein, but not other *Os*GS3 variants, failed to interact with either RGA1 (Figure 6D) or any of the rice extra-large Gα subunits (*Os*XLGs; Cantos et al., 2023) (Figure S5D-G), strengthening our conclusion that the truncated *Os*GS3-3 protein is non-functional. We similarly show that the *Os*GS3-3 protein is unable to dimerize with Gβ (Figure 6C), which thus eliminates *Os*GS3-3 competition for Gβ dimerization with the Gγ subunits that promote grain elongation, *Os*DEP1 and *Os*GGC2 (Sun et al., 2018), resulting in a long-grain phenotype. Contrastingly, multiple *Os*GS3 mutants that result in a super short-grain phenotype (Takano-Kai et al., 2013) retain the ability to interact with *Os*RGB1, namely, *Os*GS3-4, *Os*GS3-5, *Os*GS3-6 and *Os*GS3-7 (Figures 6B and 6C), consistent with the ability of these proteins to partially suppress *Os*DEP1 and *Os*GGC2 dimerization with Gβ, despite presumably not retaining full intrinsic signaling functionality of their own.

As our observations of genetic interaction between different Gγ subunit variants are consistent with the concept of Gγ competition for dimerization with the Gβ subunit, introgression of different Gγ alleles into elite varieties is likely to have unintended effects on signaling from other heterotrimeric G protein components. As such scenarios of pleiotropism and genetic interactions are not uncommon, we envision that Oryza CLIMtools will facilitate the identification of sources of complementary genetic variation and suitable genetic backgrounds for rational breeding and genetic improvement programs; moreover, unlike previous studies, CLIMtools provides this information in the context of environmental adaptations. As an example, we discovered that the full-length *OsGS3* allele is found in areas of lower PET – and lower temperatures – in both Japonica (Figures 4B and C) and Indica landraces (Figures 5B and C). Consistent with these results, our previous work identified a role for the Arabidopsis ortholog *AtAGG3* (Chakravorty et al., 2011) in adaptation to cold temperatures and the association of a missense (minor) variant in *AtAGG3* with colder areas in northern Eurasia (Ferrero-Serrano and Assmann, 2019).

*OsDEP1*, originally identified as an erect panicle QTL (Kong et al., 2007; Yan et al., 2007), encodes another Gγ protein that, like *OsGS3*, is a Type C Gγ subunit with a Cys-rich tail. Natural variation in *OsDEP1* regulates meristematic activity, with a previously described variant resulting in a reduced length of the inflorescence internode, erect and shorter panicles, an increased number of grains per panicle, and a consequent increase in grain yield (Huang et al., 2009). This variant involves the replacement of a 637-bp stretch of the middle of exon 5 with a 12-bp sequence, creating a premature stop codon and the loss of 230 residues from the C terminus (Huang et al., 2009). The variant we identify here is a missense variant that also occurs in the middle of the 5^th^ exon (Figure 5A). We show that, just as for *OsGS3*, variation in *OsDEP1* is also associated with coldest quarter PET (Figures 5B and 5C). However, based on our analysis of panicle length in Indica landraces, we propose that natural variation in *OsDEP1*, but not *OsGS3*, is responsible for the variation in this phenotype in Indica varieties. We reach this conclusion based on the association of the minor *OsDEP1* allele with shorter panicle phenotypes and the major *OsDEP1* allele with longer panicle phenotypes, independent of a full-length/truncated *OsGS3* allele (Figure 5E). While this conclusion might be challenged by our observation of differences in panicle length in Japonica landraces associated with natural variation in *OsRGA1* and *Os*GS3 (Figure S4), we suggest that this apparent discrepancy may be due to an interaction with the *OsRGA1* allele in Japonica that is not possible in Indica, which completely lacks the variants in *OsRGA1* that we describe. Alternatively, there may be a genetic interaction with the different haplotypic versions of *OsGCC2* present in Indica vs. Japonica (Figure 6A).

We recently proposed a mechanism regulated by G proteins, particularly the Gα subunit, that involves a trade-off between growth and stress tolerance that is critical to mediate adaptability (Ferrero-Serrano and Chakravorty, 2023). We hypothesize that landraces from areas with lower coldest quarter PET (typically, at higher latitudes; Figure 3C), experience a “less predictable” environment (Figure 3A) and trade-off their stress tolerance with a derived cost in growth that is illustrated by the reduced stature associated with the minor *OsRGA1* allele (Figure 4D and E). This point is illustrated not only by the increased drought tolerance of the dwarf *d1* mutant, but also by previous findings relating loss-of-function mutants of Gα with resistance to diverse stresses in other species (Zhang et al., 2008; Nilson and Assmann, 2010; Jangam et al., 2016; Ferrero-Serrano et al., 2018; Cui et al., 2020). We also highlight the association of natural variation in the Gγ subunit with the same environmental gradient we found in association with *OsRGA1* variants, PET during the coldest quarter, and in this case, also with reproductive traits that are important to maximize yield (Figure 6E). These results highlight the potential of modulating G protein function for vegetative stress tolerance, and the associated interplay between stress tolerance, growth, and yield.

## Conclusion

There is a mismatch between the current distributions of crop varieties and climate suitability for their production. Breeding programs that introduce genetic diversity from local landraces can help to address this challenge (Hoisington et al., 1999; Fu and Somers, 2009; Zhang et al., 2017; Renard and Tilman, 2019) but will benefit from taking environmental factors into account.

We have demonstrated the utility of Oryza CLIMtools to uncover adaptive genetic variation that is associated with existing natural variation in agronomic traits. The results described in this report highlight the promise of introducing natural genetic variation from landraces into modern rice varieties. The introduction of naturally occurring genetic variation into breeding programs is not new, but the assessment of genotypic interaction with the local environment adds a new dimension to approaches to create and identify varieties that are better adapted to their local climate, and foster a more sustainable food production system.

The ultimate goal is to incorporate the information derived from landraces into modern varieties and test them in appropriate multi-year and multi-environment trials, using the appropriate guidelines (Khaipho-Burch et al. 2023), to develop better-adapted varieties that will improve food security and sustainability.’

## Acknowledgments

Supported by National Institute of General Medical Sciences of the NIH under award number 5R01GM126079 to SMA and NSF-IOS-2122357 to Prof. Philip C. Bevilacqua and SMA. Kobie Kirven acknowledges support from NIH training grant 5T32GM102057. The content is solely the responsibility of the authors and does not necessarily represent the official views of the NIH. We thank Prof. D. Ware, Dr. A. Olson, and Prof. P. C. Bevilacqua for helpful discussions, and comments on this manuscript. We also thank the Gramene Team (https://www.gramene.org/) for hosting Oryza CLIMtools. The Pennsylvania State University has a patent (US9434957B2), authored by SMA, AFS, and DC, on the manipulation of RGA1 activity to engineer drought tolerance and improved seed production in plants.

## Materials and Methods

### Extraction of environmental variables

We compiled a comprehensive database of environmental data for 658 Indica and 283 Japonica georeferenced landrace varieties included in the 3K Rice Genome Project (Gutaker et al., 2020) that includes 413 geo-climatic variables describing the local environment at each collection site (Figure 1, S1, and Table S1). We used the Raster package in R (Hijmans et al., 2013) to extract the geospatial raster data for the environmental conditions prevailing at each landrace’s collection site. A detailed list of relevant literature references that describe the climate dataset from which we curated the environmental information, and summary statistics for each of these 413 environmental variables can be found in Table S2.

### OryzaCLIM

We developed OryzaCLIM, a user-friendly Shiny component (Chang et al., 2015a) of Oryza CLIMtools that enables the analysis of local environmental data for each georeferenced accession (Gutaker et al., 2020) sequenced as part of the 3,000 (3K) Rice Genome Project (Wang et al., 2018). OryzaCLIM incorporates 413 geo-environmental variables that have been codified to facilitate the examination of environmental conditions at the collection sites of 658 Indica and 283 Japonica georeferenced landraces. These data points are represented on an interactive world map that allows users to analyze pairwise environmental conditions for these 941 landrace varieties.

By clicking on any of the data points on the map, users can access information about the selected variety, including its curation details and the value of the chosen environmental variable. OryzaCLIM is designed to be user-friendly and accessible, and the source code and data used to create this tool are freely available for download at GitHub (https://github.com/CLIMtools/OryzaCLIM). The code is licensed under Apache 2.0.

### GWA analysis

We obtained SNP and INDEL data from the 3K genomes (https://snp-seek.irri.org/_download.zul;jsessionid=0E0F5BCFB0EA14EBB926BEBC7B26D6E0) and converted the data from Plink format to VCF format using PLINK 2.0 (Chang et al., 2015b). We then used VCFtools (Danecek et al., 2011) to subset the resulting VCFs into separate files for 658 Indica and 283 Japonica landrace varieties, which were the focus of this study. The data were filtered to select genetic variants with less than 10% missing data and minimum allele frequency (MAF) ≥ 5%. After this initial filtering, imputation was conducted using BEAGLE v5.4 with default parameters (Browning et al., 2018).

Rice includes two major sub-species: *Oryza sativa ssp. Indica* and *Oryza sativa ssp. Japonica*. Differences between subspecies are apparent in a number of physiological and morphological traits, genetic variation, and geographical distribution (Khush, 1997; Garris et al., 2005; Huang et al., 2012; Wang et al., 2018). Our own analysis of previously characterized agronomic traits (Wang et al., 2020) in the set of landrace varieties used in this study revealed that Indica landrace varieties typically display more culms that are longer, longer leaves that are narrower, shorter stature at the seedling stage, earlier flowering, more panicles, and longer grain than Japonica varieties (Figure S6). Given such marked subspecies differences, we conducted GWA analysis separately in the Indica and Japonica subpopulations.

Different approaches have been used previously to study genotype–environment associations, including mixed models (Lasky et al. 2014; Coop et al., 2010), generalized linear models (GLM; Luo et al. 2021), and non-parametric approaches (Hancock et al. 2011; Pluess et al. 2016). We used TASSEL 5.0 (Bradbury et al., 2007) for our GWA analysis. After model evaluation, we utilized a General Linear Model (GLM) containing a correction for population structure using the first five components as the population structure matrix (GLM+PCA; Zhao et al., 2018; Figure S8; Document S2).

Using the ‘stats’ package in R (R Core Team), we calculated the Benjamini-Hochberg (Benjamini and Hochberg, 1995) and Bonferroni (Holm, 1979) values for genome-wide significance to evaluate significance thresholds for multiple tests, for each G by E association in our GWA analysis.

To predict the impact of every genetic variant, we used SnpEff (Cingolani et al., 2012) with the gene model annotation from the Ensembl genome release 53 (https://ftp.ensemblgenomes.ebi.ac.uk/pub/plants/release-53/). We focus on SNPs within transcriptional units, including introns, untranslated regions at 5’ and 3’ UTRs, and 1 kb promoter regions upstream of the most distal transcription start site. We used the “consensus transcript,” as described by SnpEff, as the variant with either the longest coding sequence (CDS) if the gene has translated transcripts or the longest cDNA (Hubbard et al., 2009; Cingolani et al., 2012). We defined ‘common variants’ as those with MAFs ≥ 5% to restrict our analysis to genetic variation that is more likely to be adaptive (Sawyer and Hartl, 1992; Ohta, 2013).

Genomic signatures of selection were calculated using VCFtools (Danecek et al., 2011). We obtained the fixation index, Fst (Wright, 1965; Weir and Cockerham, 1984), on a per-site basis and the genetic group based on Indica vs. Japonica cultivars. We also calculated nucleotide diversity (*π*) and Tajima’s *D* in 1 kb windows across the genome (Danecek et al., 2011).

### RiboSNitch prediction

A riboSNitch is a single nucleotide polymorphism (SNP) that alters the secondary structure of an RNA. To identify riboSNitches in the rice transcriptome, we used the SNPfold software tool (Halvorsen et al., 2010). SNPfold predicts riboSNitches by using a thermodynamic model that calculates the partition function and base pairing probability matrices for the reference and variant sequences defined by any given SNP. SNPfold then compares the column sums of the base-pairing probability matrices by computing the Pearson correlation coefficient. A low SNPfold score, and thus a low Pearson correlation coefficient, indicates a potential structural change in the RNA, i.e. a riboSNitch. We suggest that SNPs are candidate riboSNitches if the correlation coefficient comparing the reference and variant sequences is less than 0.8 (Halvorsen et al., 2010), but in Oryza CLIMtools we provide the specific SNPfold score (correlation coefficient) for each variant that was predicted, so that users can choose their own threshold for designating a SNP as a riboSNtich.

SNPfold was applied to predict the presence of riboSNitches within the 3,914,483 natural genetic variants among the 658 Indica landrace varieties and 2,835,066 genetic variants among the 283 Japonica landrace varieties studied here. Each SNP in these transcriptomes was placed in the context of its surrounding 80 nucleotides (40 nucleotides upstream and 40 nucleotides downstream) and passed through the SNPfold program to determine the impact of the mutation on the local RNA structure. For Oryza CLIMtools, correlation coefficients are provided for all SNPs within annotated transcribed regions unless the SNP is located less than 40 nucleotides from a transcript end, in which case the correlation coefficient was not calculated. SNPs located less than 40 nucleotides from either end of a transcript are designated as Not Applicable in Oryza CLIMtools.

### Oryza GenoCLIM

Oryza GenoCLIM is a user-friendly Shiny (Chang et al., 2015a) component of Oryza CLIMtools. GenoCLIM is a searchable database generated with the DT package (Xie, 2016) that provides information for any locus ID for genetic variants found to be significantly associated (FDR < 0.01) with each of the 413 geo-environmental variables included in this study. FDR thresholds are calculated for each individual environmental variable. This tool allows the user to search for a locus identifier of any given gene, annotated using the Rice Annotation Project (RAP annotation; Sakai et al., 2013), and retrieve, if present, any association between the genetic variation (SNPs and INDELs) within that gene and any given environmental variable(s) described in this study.

The database provides climate associations of genetic variants, and the user can query their ‘gene of interest’ within the Indica or Japonica varieties in an interactive database. The results shown can be filtered according to different FDR thresholds and sorted based on their association score or on any of the other population genetic indicators provided for these associations. Additionally, Oryza GenoCLIM provides a plot generated by Plotly (Sievert et al., 2018), which allows users to visualize the results interactively.

For reproducibility, the source code and data used in GenoCLIM are freely available to download on GitHub (https://github.com/CLIMtools/Oryza_GenoCLIM), with the code licensed under Apache 2.0.

### Oryza CLIMGeno

Oryza CLIMGeno comprises a software suite of Shiny apps (Chang et al., 2015a) to analyze the relationships between genetic variation and environmental factors in rice. CLIMGeno is built on Zbrowse (Ziegler et al., 2015), a genome browser tool. Using CLIMGeno, users can visualize the top 2,500 genetic variants (SNPs and INDELS) associated with each of the 413 environmental variables analyzed in this study. Specifically, the tool displays the highest-scoring variants in Indica (658) or Japonica (283) rice varieties.

In Oryza CLIMGeno, the ‘manage’ tab provides users with a dropdown menu to select an environmental variable of interest. Once selected, the tool automatically generates a list of the 2,500 genetic variants (SNPs and INDELs) with the strongest association score that have an FDR < 0.01 for that variable. Users can navigate to the ‘data table’ tab to select and sort genetic variation based on various details such as position, allele frequency, locus identification, locus description, riboSNitch prediction, or SNP effect.

The ‘whole-genome view’ tab displays an interactive Manhattan plot that represents the genetic variation associated with the geo-climatic variable previously selected by the ‘whole-genome view’ tab. The x-axis shows each of the twelve rice chromosomes, while the y-axis displays the score (negative logarithm of the p-value), depicting the strength of association between genetic variability and any given environmental variable. Users can then filter the genetic variants to display based on their predicted effect using the ‘select effect’ box. Hovering over the data points provides information on the genetic effect of a given genetic variant.

Clicking on a genetic variant takes users to the ‘chromosome view’ tab, which displays two Manhattan plots corresponding to the chromosome where the selected variant is located. The top plot allows users to browse genetic variation on that chromosome and zoom in on any given region. In contrast, the bottom plot combines a Manhattan plot with an interactive display and annotation of genes found in the region of interest according to the gene model annotation from the Ensembl genome release (https://ftp.ensemblgenomes.ebi.ac.uk/pub/plants/release-53/). Users can explore the region of the chromosome surrounding the genetic variant(s) of interest by selecting a window size or using the menu on the left. Lastly, the ‘annotations table’ tab provides an interactive table with annotation information on the region of interest selected on the Manhattan plot, complemented by a graphical annotation display.

For reproducibility, the source code and data used in Oryza CLIMGeno for both Indica and Japonica varieties are freely available to download at GitHub (https://github.com/CLIMtools/Oryza_CLIMGeno_Indica and https://github.com/CLIMtools/Oryza_CLIMGeno_Japonica), with code licensed under Apache 2.0.

### Plant growth conditions and physiological characterization of drought tolerance in an *OsRGA1* mutant, *d1*

Wild-type (WT) and *d1* mutants were grown in a greenhouse in two-gallon pots containing Metro-mix 360 potting mixture. Greenhouse temperatures were maintained at 30°C during the day and 20°C during the night in a 16:8 day:night cycle with light supplied as natural daylight supplemented with 1000W metal halide lamps (Philips Lighting Co., Somerset, NJ) for the duration of the light cycle. Light intensity was ∼500 μmol m^-2^ s^-1^ PPFD. Plants were maintained under well-watered conditions until 60 days after emergence. At that point, we introduced two different drought treatments, which resulted in moderate and severe stress under the chosen treatments. Plants were watered twice daily to maintain at all times, by lysimetry, the three water treatments: well-watered/no drought (ND, 100% soil relative water content), moderate drought (MD, 45% SRWC) and severe drought (SD, 30% SRWC). There were 7 replicates per water treatment per genotype, for a total of 42 plants. Relative soil water content was kept constant from the start of the drought treatment until seed maturity, by continued twice daily watering to the appropriate weight and continuous monitoring with a Campbell Scientific TDR 100 system (Campbell Scientific Inc., Logan, Utah, USA) with a custom-made probe of 20 cm length. It was determined that in-between watering events, the relative soil water content of the soil did not vary more than 5%.

For gas exchange measurements, regions of flag leaves were enclosed in the chamber of a portable gas exchange system (LI-COR 6400 IRGA with an integrated 6400-40 leaf chamber fluorometer, Li-COR, Inc., Lincoln, NE, USA). Leaves were measured at the point of maximal width. The leaf chamber temperature was kept constant at 30°C. Airflow in the chamber was adjusted to 300 μmol s^-1^. Steady-state measurements of the different physiological parameters were performed at 500 μmol m^-2^ s^-1^ PPFD provided by the Li-COR red/blue LED system with blue light accounting for 10% of the total photon flux.

### Amplification of G protein subunits

The G protein subunits of rice are: *OsRGA1* (Os05g0333200), *OsXLG1* (Os12g0593000), *OsXLG3a* (Os11g0206700), *OsXLG3b* (Os06g0111400), *OsXLG4* (Os10g0117800), *OsRGB1* (Os03g0669200), *OsRGG1* (Os03g0635100), *OsRGG2* (Os02g0137800)*, OsGS3* (Os03g0407400), *OsDEP1* (Os09g0441900) and *OsGGC2* (Os08g0456600). G protein subunit open reading frames were amplified from seedling or flower cDNA of the Japonica Taichung 65 variety and cloned into the pCR™8/GW/TOPO™ entry vector (Thermo). The 5’ end of the *OsXLG4* ORF includes partially repetitive regions of 80+ % GC content and could not be amplified from cDNA. To circumvent this issue, the region of the *OsXLG4* cDNA encoding amino acids 198-828 was amplified from Taichung 65 cDNA and joined by overlap extension PCR to a gBlock fragment (IDT) of the region encoding amino acids 1-197 that was designed to encode the correct amino acids with reduced GC content. Overlap extension PCR was used to generate deletions of *OsGS3* previously described elsewhere (*OsGS3-4*, *OsGS3-5*, *OsGS3-7*), while an *OsGS3-3* clone was created by incorporating the C55* mutation (chromosome 3, position 16733441) into a 3’ primer that only amplifies the shortened open reading frame of the *OsGS3-3* allele. *OsGS3-6*, *Osdep1* and *OsDn1-1* was cloned by a similar method to *OsGS3-3*, see Table S3 for primers used. The *OsGS3-3* and *OsGS3-4* alleles were originally named by Mao et al. (Mao et al., 2010). We continued this naming convention with the alleles described by Takano-Kai et al. (2013), i.e. *OsGS3-5* corresponds to the allele identified in the Podiwi-A8 cultivar, *OsGS3-6* to the allele from JC73-4, JC157, and ABRI, *OsGS3-7* to the allele from H343 and Tumo-Tumo, and the allele identified in ARC7291 and TAL214 cultivars correspond to the already described *OsGS3-4* allele (Mao et al., 2010) identified in the Chuan 7 cultivar (Fan et al., 2006). *Osdep1* was described by Huang *et al*. (Huang et al., 2009), and *OsDn1-1* by Taguchi-Shiobara *et al*. (Taguchi-Shiobara et al., 2011). Multiple alignments of the protein sequences of the above *OsGS3* and *OsDEP1* alleles were conducted using Clustal Omega (Nguyen et al., 2016).

### Yeast protein-protein interaction assays

Sequence-verified Gateway entry clones were mobilized into pDEST-GADT7 and pDEST-GBKT7 destination yeast 2-hybrid vectors (Uetz et al., 2006) by Gateway LR recombination. For yeast 3-hybrid, in which the interaction of two proteins is assayed by a yeast 2-hybrid approach in the presence of a third “bridge” protein, *OsRGB1* was amplified with flanking NotI and BamHI sites to clone into the NotI/BglII sites of pBridge MCS2 (Takara) as the “bridge” protein. Gγ subunits were amplified with flanking BglII sites to ligate into the compatible BamHI site (*OsGS3* alleles), or EcoRI and SalI restriction sites (*OsRGG1*, *OsRGG2*, *OsGGC2*), or EcoRI and BamHI restriction sites (*OsDEP1* alleles) to clone into MCS1 of pBridge as GAL4BD fusions. Yeast 2-/3-hybrid assays were performed as previously described (Chakravorty et al., 2015).

