## Supplementary material for "Oryza CLIMtools: A Genome-Environment Association Resource Reveals Adaptive Roles for Heterotrimeric G Proteins in the Regulation of Rice Agronomic Traits": Table S3 and SI Figures

**Table S3. Primer sequences used to generate clones of heterotrimeric G protein subunits and alternative alleles. Within primer names 'f' corresponds to forward primers, 'rs' corresponds to reverse primers containing a stop codon and lower case 'o' indicates a primer used for overlap-extension PCR.**

| Primer | Sequence | Use |
|---|---|---|
| **OsRGA1f** | ATGGGCTCATCCTGTAGC | Cloning of *OsRGA1* |
| **OsRGA1rs** | TCAAGTTCCTTCCCTGGAG | |
| **OsXLG1f** | ATGGCGGCGGCGGGGGAGTACTCGTTCG | Cloning of *OsXLG1* |
| **OsXLG1rs** | TCAGTGAGAGTAGGAGCTCGGCTCCTCG | |
| **OsXLG3af** | ATGGCGGAGGCGGTGGATGGGGGCAGC | Cloning of *OsXLG3a* |
| **OsXLG3ars** | TCATTCTGCTCGGATGAAAGGAGAGGAGC | |
| **OsXLG3bf** | ATGGCCGGAGCGGGAGCCGTGGGGG | Cloning of *OsXLG3b* |
| **OsXLG3brs** | TCATTCTTGTCTGATCAGACGTGAGGAGC | |
| **OsXLG4f** | ATGGCGTCCTCACCTCCCGCTCCGCC | Cloning of 5' vs 3' *OsXLG4* fragments followed by overlap-extension PCR |
| **OsXLG4R1** | CTTGCGGCCCTCCGGCATGGACCCCAT | |
| **OsXLG4F1** | ATGGGGTCCATGCCGGAGGGCCGCAAG | |
| **OsXLG4rs** | TTAGGATGGAGAGGTCGTCATGTCCACACTGTACGAG | |
| **OsRGB1f** | ATGGCGTCCGTGGCGGAGCTCAAGGAGAAGC | Cloning of *OsRGB1* |
| **OsRGB1rs** | TCAAACTATTTTCCGGTGTCCGCTGAAGGCCCAAATC | |
| **OsRGG1f** | ATGCAGGCCGGAGGAGGAGGGGACG | Cloning of *OsRGG1* |
| **OsRGG1rs** | TCACAAAAACCAGCATTTGCATCTGCGCAG | |
| **OsRGG2f** | ATGAGGGGGGAGGCGAACGGGGAGG | Cloning of *OsRGG2* |
| **OsRGG2rs** | CTAGGAAAAATCTGAGCCTTTGGATGCCCACC | |
| **OsGS3f** | ATGGCAATGGCGGCGGCGCCCCGGCCC | Cloning of *OsGS3* |
| **OsGS3rs** | TCACAAGCAGGGGGGGCAGCAACGAGGGAC | |
| **OsGS3-3rs** | TCAGCATCTGGAGGCAGC | Generation of *OsGS3* alleles |
| **OsGS3-4of** | CTGCTCATCCTCTCCTCCTCCTTCAACCTCAAGAGG | |
| **OsGS3-4or** | TGAGGTTGAAGGAGGAGGAGAGGATGAGCAGCCGCCGGC | |
| **OsGS3-5of** | GCAGAGCAAGTCGTGCTGCCTCAGCTACCTC | |
| **OsGS3-5or** | GAGGCAGCACGACTTGCTCTGCACAAACAGCG | |
| **OsGS3-6rs** | TTAAGTACGCGCGCAGCAGCAGCTCGGCCTCT | |
| **OsGS3-7of** | TGCTGCAGCAGCGCTCATCCTCCTCCTCCTCCTTCA | |
| **OsGS3-7or** | TGAAGGAGGAGGAGGAGGAGGATGAGCGCTGCTGCAGCAGCAGA | |
| **OsDEP1f** | ATGGGGGAGGAGGCGGTGGTGATGGAGGC | Cloning of *OsDEP1* |
| **OsDEP1rs** | TCAACATAAGCAACCACTGAGACAGCATGGGTTACGACACCGTGGG | |
| **Osdep1rs** | CTAGATGTTGAAGCAGGTGCA | Generation of *OsDEP1* alleles |
| **OsDn1-1rs** | CTAGCAGTTCGGTTTGCAGC | |
| **OsGGC2f** | ATGGGGGAGGCGCCGAGGCCC | Cloning of *OsGGC2* |
| **OsGGC2rs** | TCAGCATAAGCATCCGCCGGCACAG | |

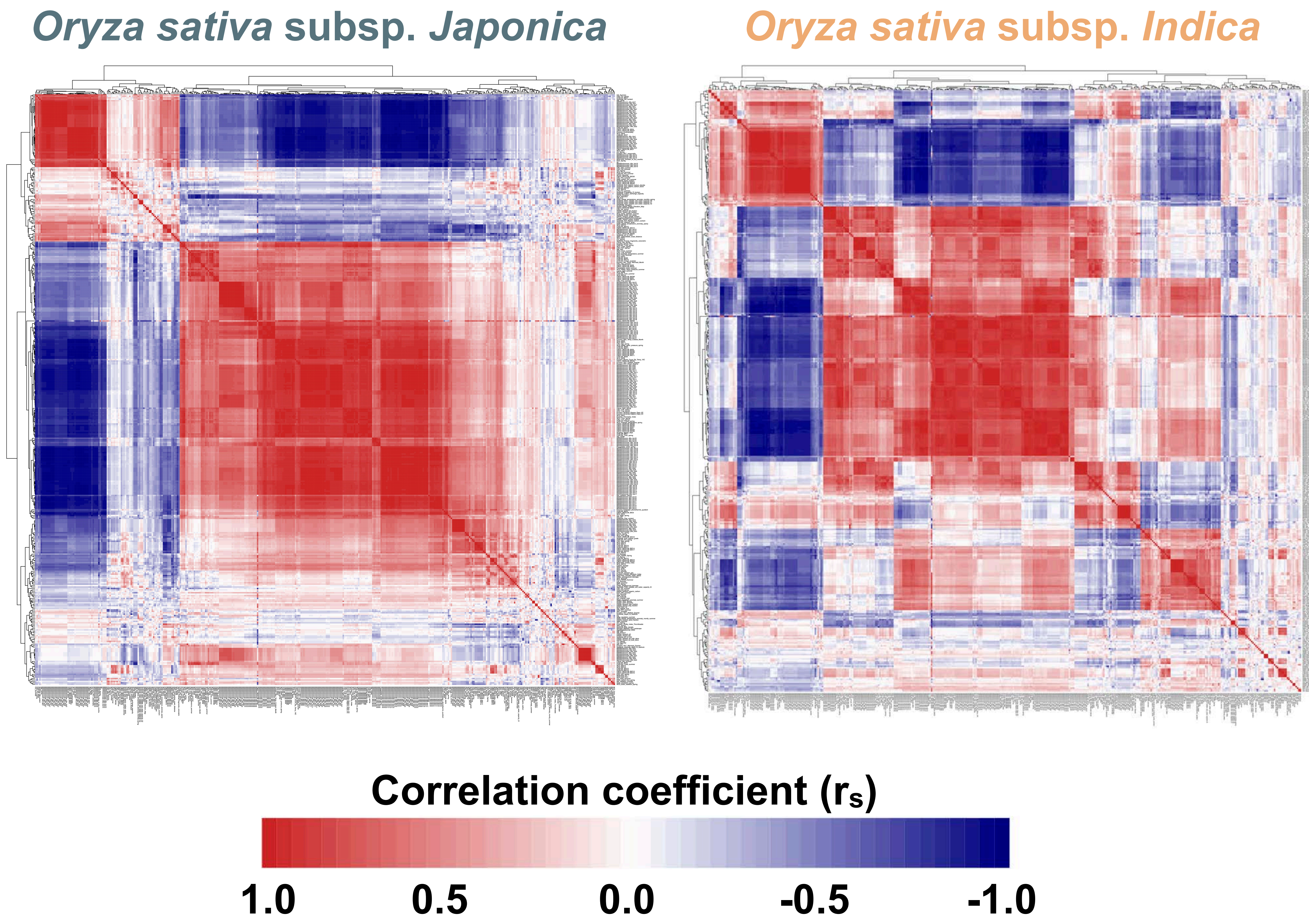


**Figure S1. Correlations among the environmental variables presented in this study for the set of 413 geo-environmental variables extracted for the local environment of the 658 Indica and 283 Japonica georeferenced landrace varieties depicted in this study.** The environmental variables in this plot are ordered by hierarchical clustering. White indicates a lack of correlation, with a continuous color scale ranging from dark blue (strong positive correlation) to dark red (strong negative correlation).

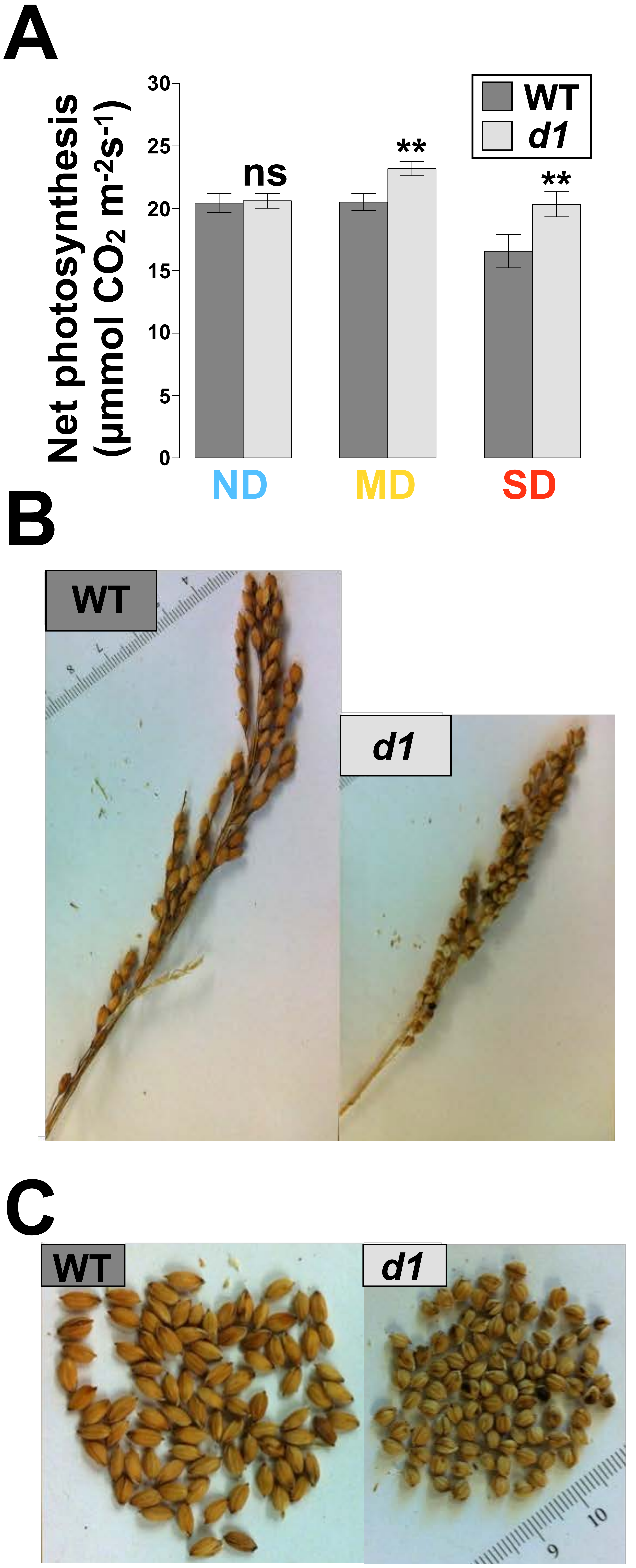


**Figure S2. *OsRGA1* loss of function mutant *d1* displays greater photosynthesis under drought and distinctly shorter panicles and seeds than wild-type (Taichung 65). A.** In the absence of drought (ND), *d1* displays similar photosynthetic rates relative to wild-type (WT), but under moderate drought (MD; 45% relative soil water content) and severe drought (SD; 35 % relative soil water content), *d1* photosynthesis exceeds that of wild-type. **B.** *d1* plants exhibit shorter panicles and **C.** rounder and shorter seeds.

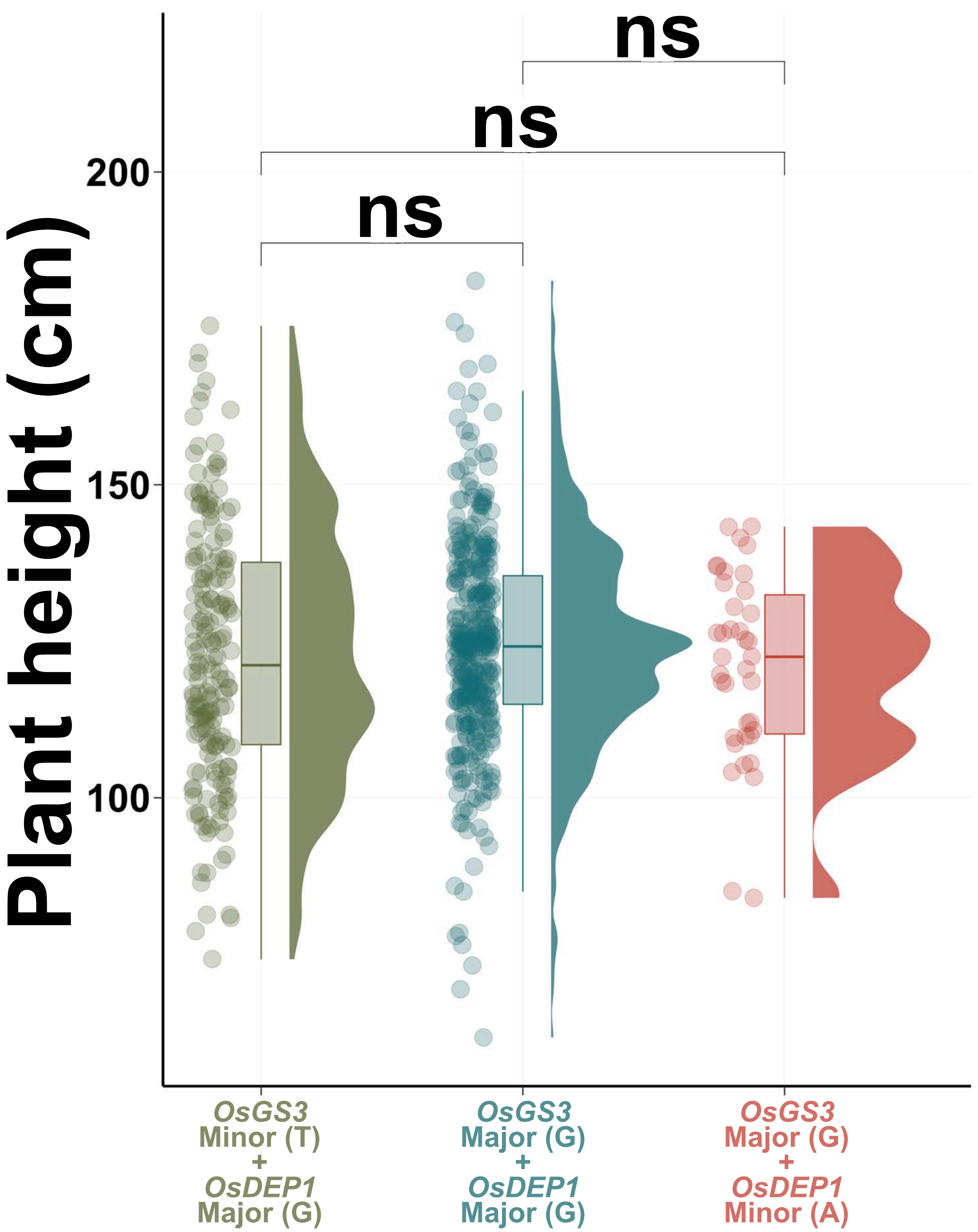


**Figure S3. Genetic variation in *OsGS3* and *OsDEP1* is not associated with differential plant height.** Raincloud plots illustrate that the presence of *OsGS3* and *OsDEP1* variants in Indica varieties is not associated with differential plant height.

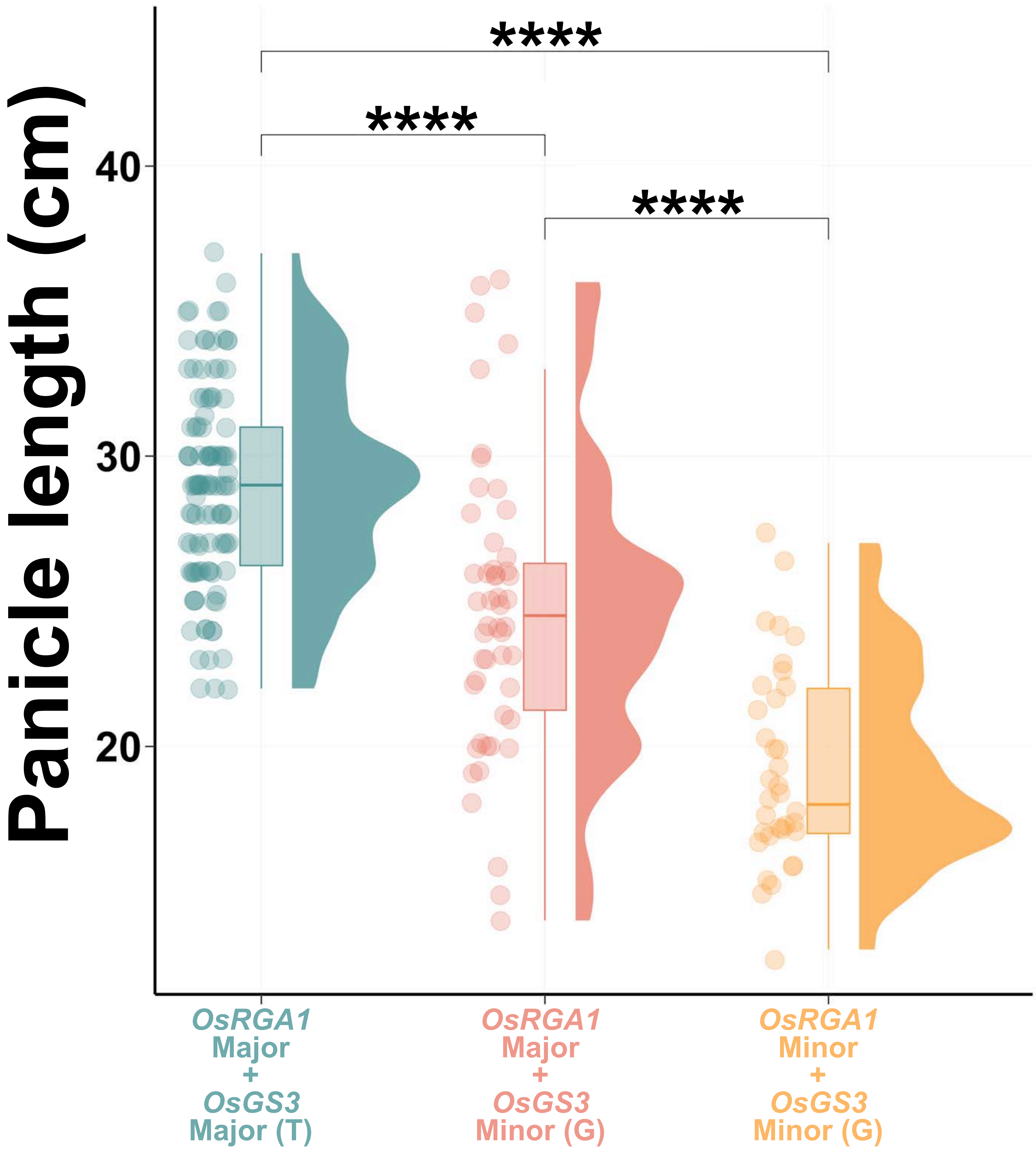


**Figure S4. Genetic variation in *OsRGA1* and *OsGS3* is associated with differences in panicle length in Japonica landraces.** Raincloud plots illustrate that the presence of *OsRGA1* and *OsGS3* variants in Japonica landraces is associated with differential panicle length. Four asterisks (****) indicate a *p-value* < 0.0001 using the non-parametric Wilcoxon test.

# A

```
GS3       MAMAAAPRPKSPPAPPDPCGRHRLQLAVDALHREIGFLEGEINSIEGIHAASRCCREVDEFIGRTPDPFITISSEKRSHDHSHHFLKKFRCLCRASACCL   100
GS3-3     MAMAAAPRPKSPPAPPDPCGRHRLQLAVDALHREIGFLEGEINSIEGIHAASRC*---------------------------------------------   54
GS3-4     MAMAAAPRPKSPPAPPDPCGRHRLQLAVDALHREIGFLEGEINSIEGIHAASRCCREVDEFIGRTPDPFITISSEKRSHDHSHHFLKKFRCLCRASACCL   100
GS3-5     MAMAAAPRPKSPPAPPDPCGRHRLQLAVDALHREIGFLEGEINSIEGIHAASRCCREVDEFIGRTPDPFITISSEKRSHDHSHHFLKKFRCLCRASRAAS   100
GS3-6     MAMAAAPRPKSPPAPPDPCGRHRLQLAVDALHREIGFLEGEINSIEGIHAASRCCREVDEFIGRTPDPFITISSEKRSHDHSHHFLKKFRCLCRASACCL   100
GS3-7     MAMAAAPRPKSPPAPPDPCGRHRLQLAVDALHREIGFLEGEINSIEGIHAASRCCREVDEFIGRTPDPFITISSEKRSHDHSHHFLKKFRCLCRASACCL   100
          ******************************************************

GS3       SYLSWICCCSSAAGGCSSSSSSSFNLKRPSCCCNCNCNCCSSSSSSCGAALTKSPCRCRRRSCCCRRCCCGGVGVRACASCSCSPPCACCAPPCAGCSCR   200
GS3-3     ----------------------------------------------------------------------------------------------------   54
GS3-4     SYLSWICCCSSAAGGCSSSPPPPSTSRGRAAAATA------------TATAAPPPPHVGRR*--------------------------------------   149
GS3-5     ATSPGSAA-AAAPPAAAHPPPPPSTSRGRAAAATA-----------TATAAPPPPPHVGRR*--------------------------------------   149
GS3-6     SYLSWICCCSSAAGGCSSSSSSSFNLKRPSCCCART*---------------------------------------------------------------   136
GS3-7     SYLSWICCCSSAHP----PPPPPSTSRGRAAAATA-----------TATAAPPPPPHVGRR*--------------------------------------   146

GS3       CTCPCPCPGGCSCACPACRCCCGVPRCCPPCL*       232
GS3-3     ---------------------------------       54
GS3-4     ---------------------------------       149
GS3-5     ---------------------------------       149
GS3-6     ---------------------------------       136
GS3-7     ---------------------------------       146
```

# B

```
DEP1      MGEEAVVMEAPRPKSPPRYPDLCGRRRMQLEVQILSREITFLKDELHFLEGAQPVSRSGCIKEINEFVGTKHDPLIPTKRRRHRSCRLFRWIGSKLCICI   100
dep1      MGEEAVVMEAPRPKSPPRYPDLCGRRRMQLEVQILSREITFLKDELHFLEGAQPVSRSGCIKEINEFVGTKHDPLIPTKRRRHRSCRLFRWIGSKLCICI   100
Dn1-1     MGEEAVVMEAPRPKSPPRYPDLCGRRRMQLEVQILSREITFLKDELHFLEGAQPVSRSGCIKEINEFVGTKHDPLIPTKRRRHRSCRLFRWIGSKLCICI   100
          ****************************************************************************************************

DEP1      SCLCYCCKCSPKCKRPRCLNCSCSSCCDEPCCKPNCSACCAGSCCSPDCCSCCKPNCSCCKTPSCCKPNCSCSCPSCSSCCDTSCCKPSCTCFNIFSCFK   200
dep1      SCLCYCCKCSPKCKRPRCLNCSCSSCCDEPCCKPNCSACCAGSCCSPDCCSCCKPNCSCCKTPSCCKPNCSCSCPSCSSCCDTSCCKPSCTCFNI*----   195
Dn1-1     SCLCYCCKCSPKCKRPRCLNCSCSSCCDEPCCKPNCSACCAGSCCSPDCCSCCKPNCSCCKTPSCCKPNC*-----------------------------   170
          **********************************************************************

DEP1      SLYSCFKIPSCFKSQCNCSSPNCCTCTLPSCSCKGCACPSCGCNGCGCPSCGCNGCGCPSCGCNGCGLPSCGCNGCGSCSCAQCKPDCGSCSTNCCSCKP   300
dep1      ----------------------------------------------------------------------------------------------------   195
Dn1-1     ----------------------------------------------------------------------------------------------------   170

DEP1      SCNGCCGEQCCRCADCFSCSCPRCSSCFNIFKCSCAGCCSSLCKCPCTTQCFSCQSSCCKRQPSCCKCQSSCCEGQPSCCEGHCCSLPKPSCPECSCGCV   400
dep1      ----------------------------------------------------------------------------------------------------   195
Dn1-1     ----------------------------------------------------------------------------------------------------   170

DEP1      WSCKNCTEGCRCPRCRNPCCLSGCLC* 426
dep1      ---------------------------195
Dn1-1     ---------------------------170
```

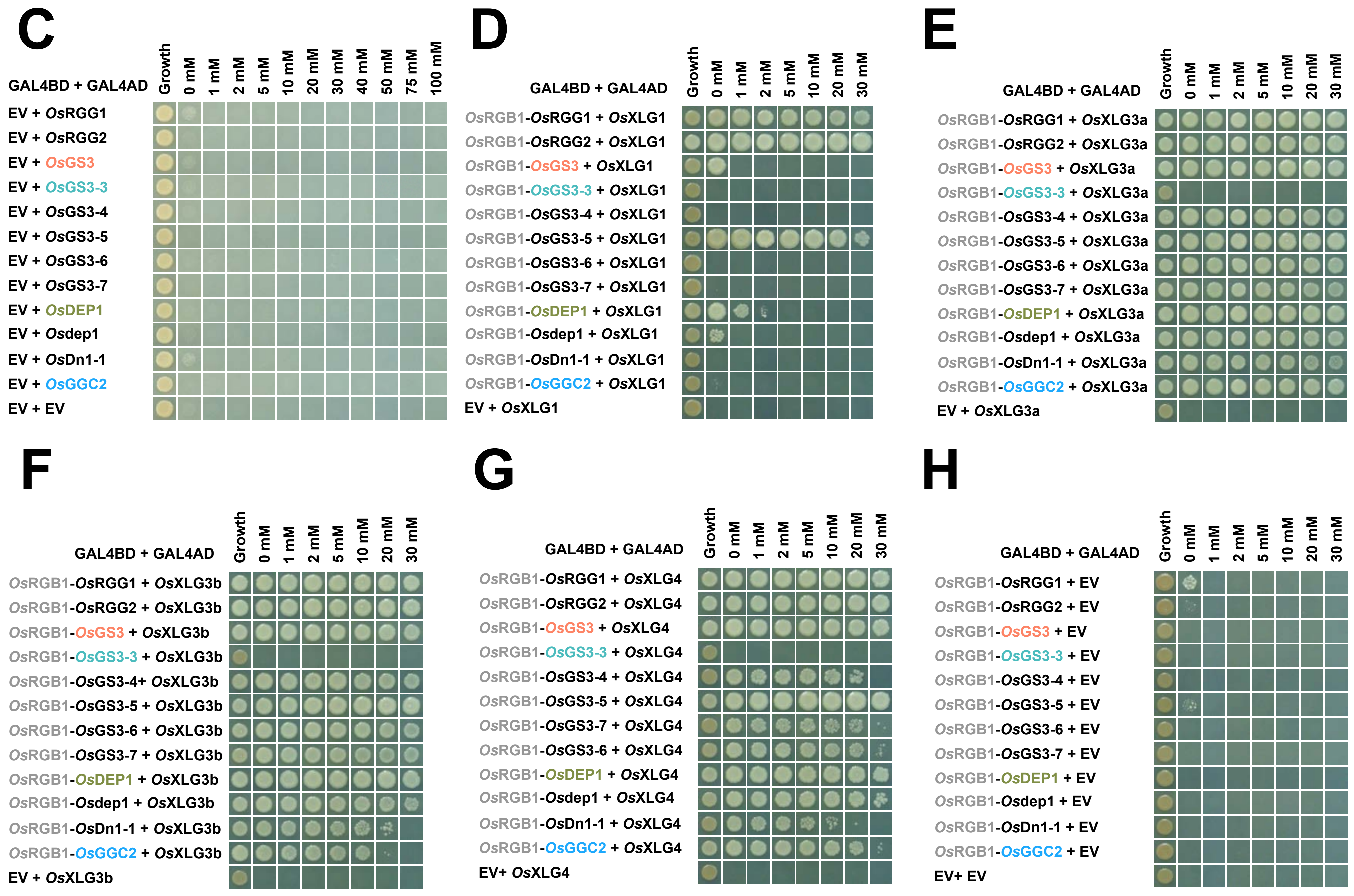


**Figure S5. *Os*GS3-3 does not interact with heterotrimeric G protein subunits.** Multiple sequence alignments of previously described naturally occurring truncations of **A.** *Os*GS3 and **B.** *Os*DEP1. **C.** Empty vector-Gγ yeast 2-hybrid panel (negative control for Figure 6C), demonstrating a lack of substantial growth and thus absence of aberrant autoactivation in the assay. Yeast 3-hybrid assays between Gβγ dimers and the extra-large Gα subunits: **D.** *Os*XLG1, **E.** *Os*XLG3a, **F.** *Os*XLG3b and **G.** *Os*XLG4. Assays were carried out in parallel with the Gβγ-*Os*RGA1 assay depicted in Figure 6D. As for the canonical Gα, *Os*RGA1, Gβγ dimers containing *Os*GS3-3 failed to interact with any *Os*XLG. All other Gβγ dimers showed strong interaction with *Os*XLG3a, *Os*XLG3b, and *Os*XLG4. Interaction with *Os*XLG1 was Gβγ-specific. Subunits are named based on the phylogeny established by Cantos et al. (2023). **H.** Gβγ-empty vector yeast 3-hybrid panel (negative control for Figures 6D, S5D, S5E, S5F and S5G). Only *Os*RGG1-*Os*RGB1 displayed weak growth on 0 mM 3-AT medium, and therefore no autoactivation was observed that would alter the interpretation of the interactions observed for the various heterotrimeric G protein combinations.

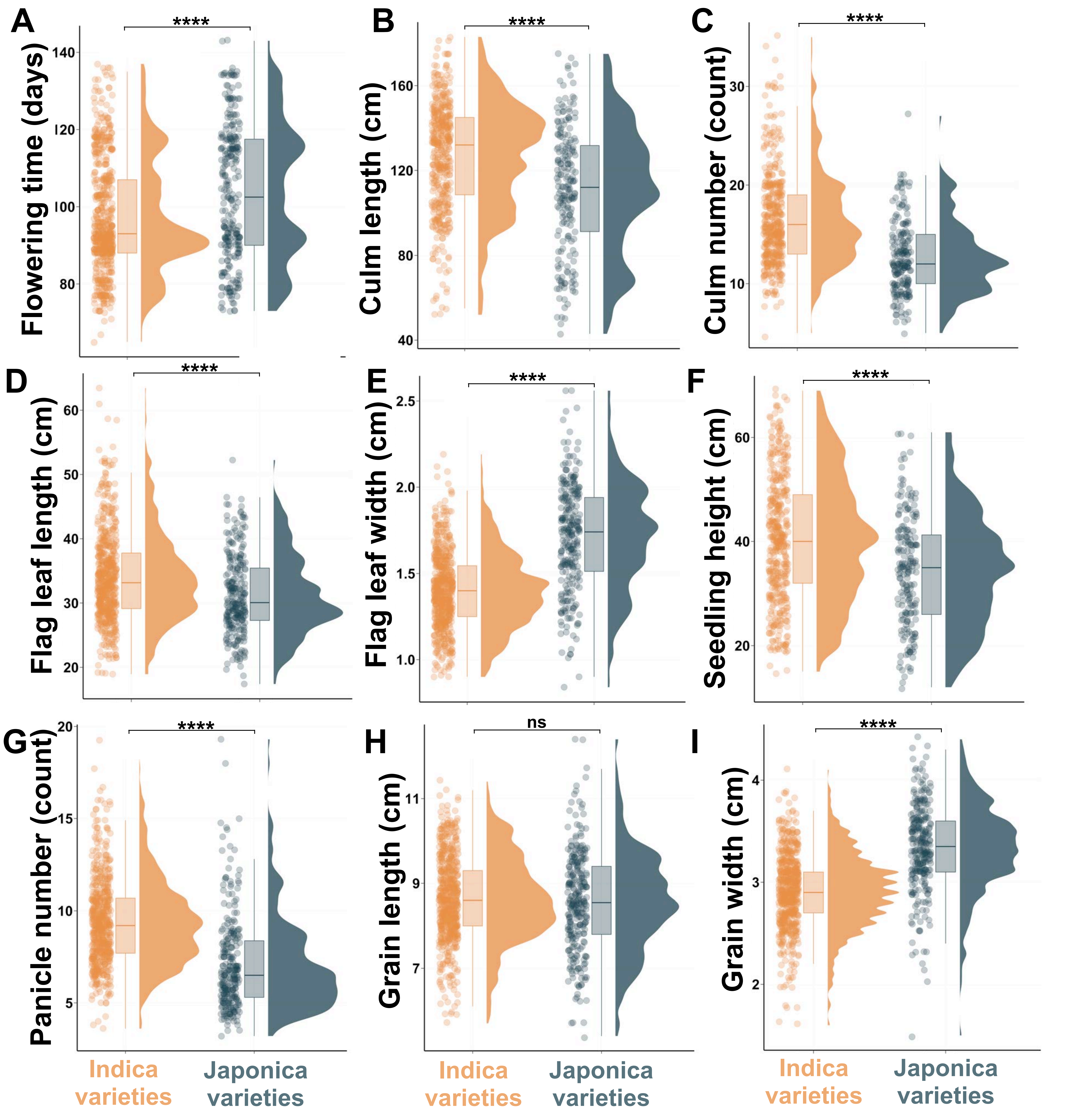


**Figure S6. The two major rice subspecies, Indica and Japonica, display distinct phenotypic characteristics.** Data corresponding to the phenotypic characterization of agronomic traits of Japonica and Indica landrace varieties were retrieved from the Functional and Genomic Breeding online resource (RFGB 2.0; Wang et al., 2020). Raincloud plots combine box plots, raw jittered data, and split-half violins. Jittered data points depict individual Indica (orange) and Japonica (navy blue) landrace varieties. In the box plots, the lower and upper boundaries indicate the 25th and 75th percentile, respectively. The whiskers below and above the box indicate the smallest value within 1.5 times the interquartile range (IQR) below the 25th percentile. The whisker above the box indicates the largest value within 1.5 times the IQR above the 75th percentile, respectively. The dark line inside the box indicates the median. Pairwise nonparametric Wilcoxon tests were conducted to assess differences between alleles between subspecies. **** ($p$ <0.0001); *** ($p$ < 0.0001 < $p$ < 0.001); ** (0.001 < $p$ < 0.01); * (0.01 < $p$ < 0.05); non-significant comparisons (ns) ($p$ > 0.05). The associations between temperature and flowering time were fitted to a linear model. Regression lines are shown in black; gray shading represents 95% confidence intervals.

A

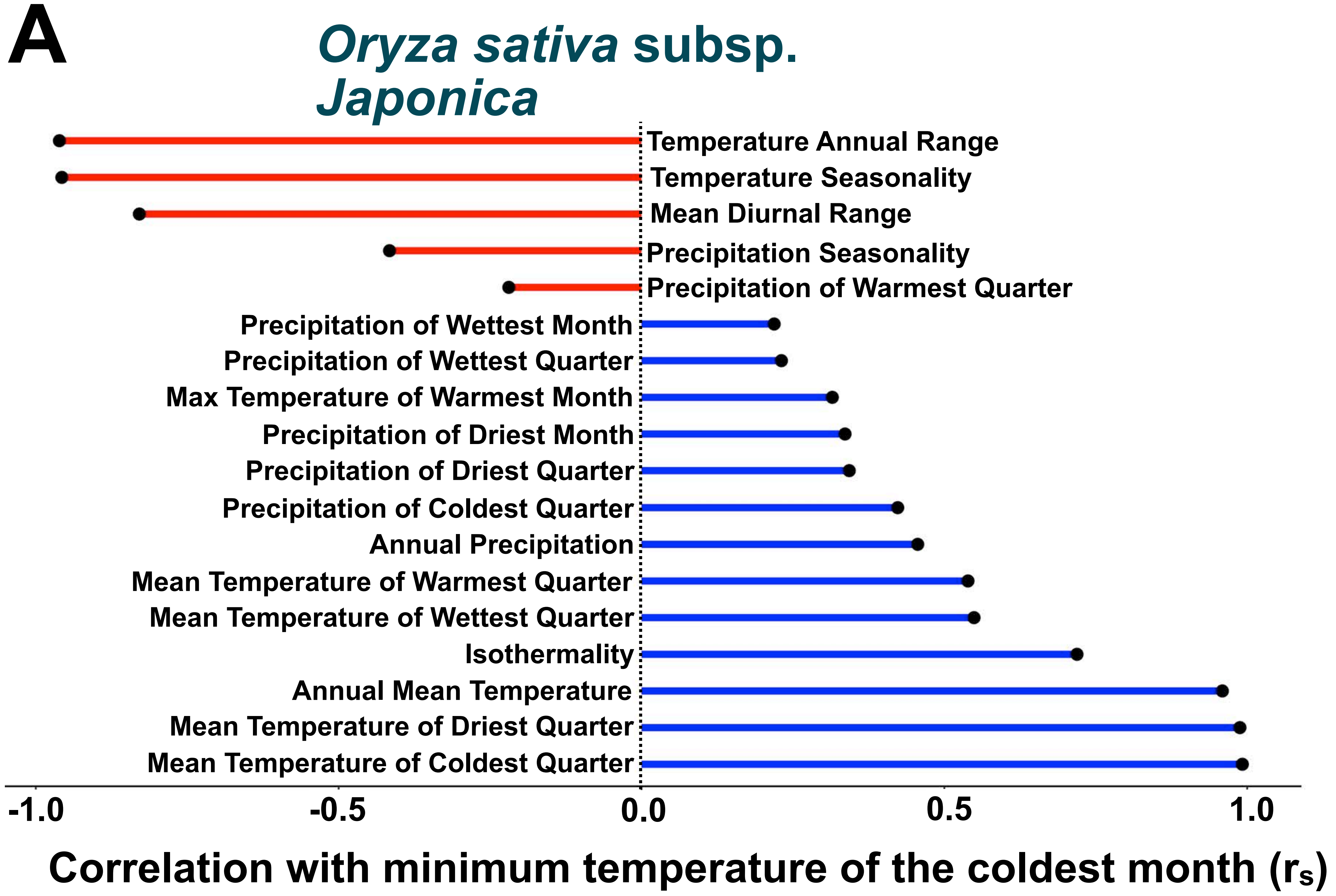


B

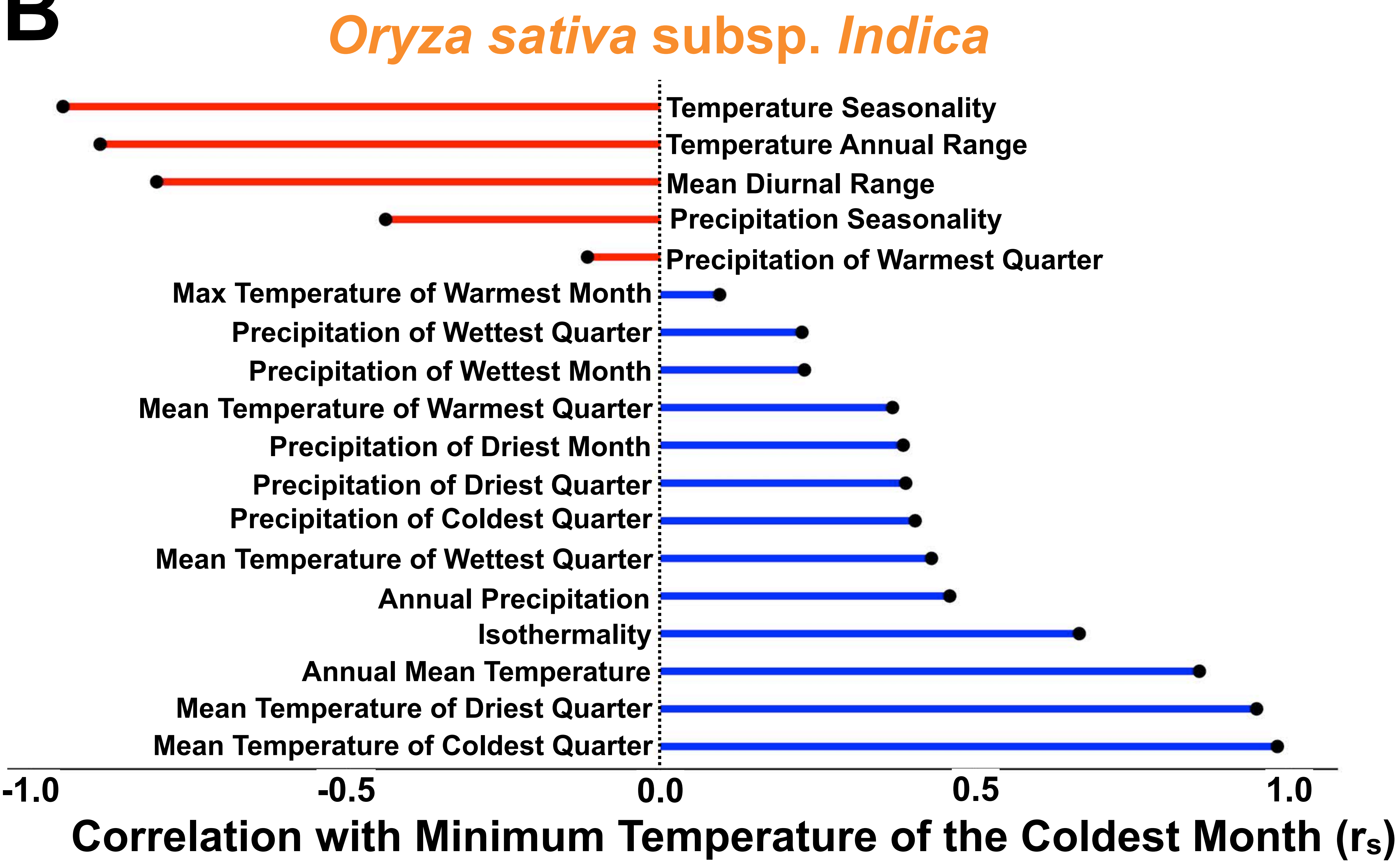


**Figure S7. The minimum temperature of the coldest month is co-associated with other climate variables.** For example, areas where both (A) Japonica and (B) Indica landraces are grown that experience colder temperatures during their coldest month also experience more temperature variability throughout the year.

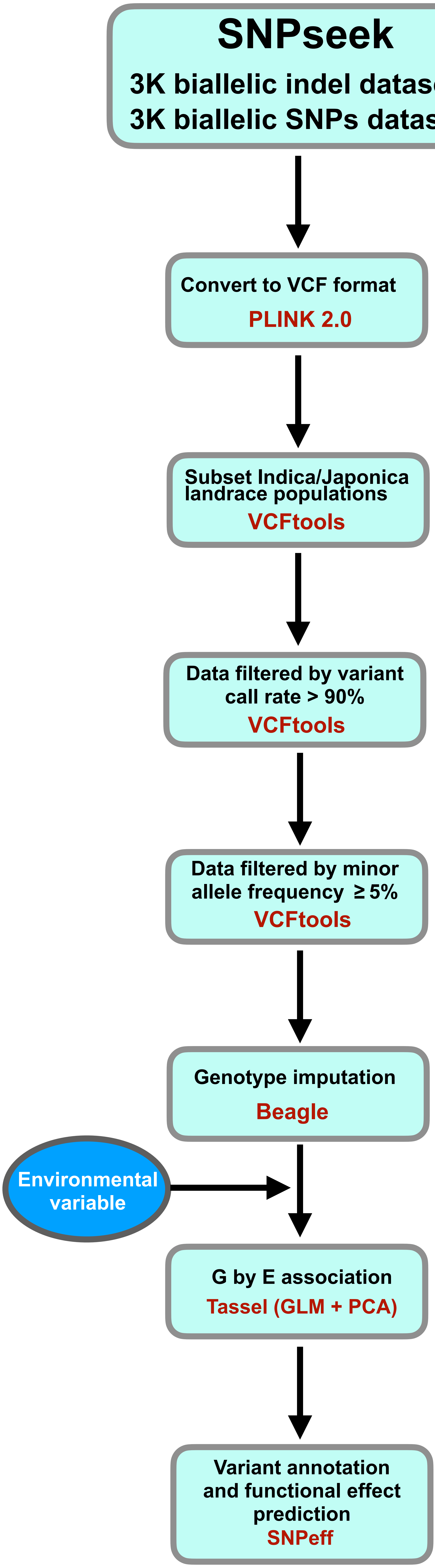


Figure S8. Flowchart of the methodology implemented to conduct genome by environment associations in this study (see Materials & Methods).
