## Supplementary material for "Oryza CLIMtools: A Genome-Environment Association Resource Reveals Adaptive Roles for Heterotrimeric G Proteins in the Regulation of Rice Agronomic Traits": Document S1

### Examples of associations with flowering time photoperiod-sensitive genes

*OsHD1* (Yano et al. 2000; Han et al. 2016; Zhang et al. 2015) is a major photoperiod sensitivity QTL in rice. Upon investigation, GenoCLIM does not return an association of *OsHD1* variants with latitude (which is a reasonable proxy for photoperiod). However, if we redo our GWA analysis using the GLM model with no correction for population structure, then six covarying *OsHD1* SNPs are found significantly associated with latitude at FDR < 0.001 (Figure 1A), consistent with a role in temperature and photoperiod seasonality. These same six covarying SNPs are also associated with flowering time (Figure 1B), based on analysis of publicly available data (Mansueto et al. 2017).

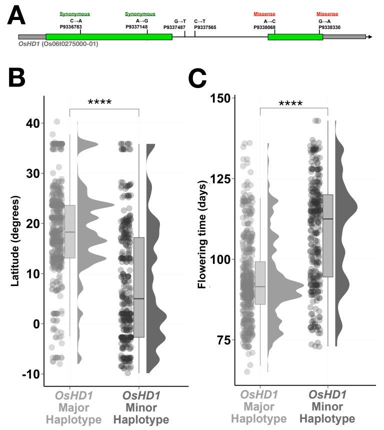


**Figure 1. Natural variation in *OsHD1* is associated with differences in heading date and latitudinal distribution, suggesting a role in photoperiod and temperature seasonality of flowering. A** We identified an allele consisting of a set of six covarying SNPs in *HEADING DATE 1* (*OsHD1*; Os06g0275000). Grey indicates UTRs, green indicates exons, and black indicates introns. The major and minor alleles of *OsHD1* are associated with: **B.** latitude, **C.** flowering time. **** (Wilcoxon, $p < 0.0001$).

A major reason for a candidate variant truly involved in a phenotype not to display the expected association following a GWA analysis, is that correction for population structure results in an increased number of false negative results (type II error) arising from over-correction. For example, as reported for African sorghum landraces, photoperiod sensitivity caused by natural variants in *MATURITY1,* the ortholog of the *OsHD2* gene that we showcase in main text Figures 2A and 2B, increases with decreasing latitude. However, this association was not significant in a GWA analysis after accounting for population structure (Lasky et al., 2015). We make this point, as well as provide other possible explanations of why a *bona fide* candidate may be missed using GWAS approaches, in the "Considerations and Limitations" document that is available at CLIMtools.

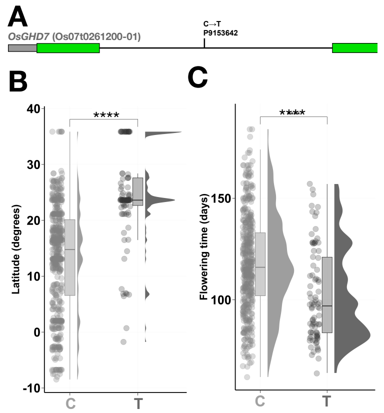


**Figure 2. Natural variation in *OsGHD7* is associated with differences in heading date and latitudinal distribution, suggesting a role in photoperiod sensitivity. A.** An intronic SNP in *OsGHD7* (Os06g0275000) that is strongly associated with: **B.** Latitude and **C.** flowering time. **** (Wilcoxon, $p < 0.0001$).

For *OsGHD7* (Yano et al. 2000; Han et al. 2016; Zhang et al. 2015), the G by E analysis provided in Oryza GenoCLIM, with correction for population structure, as is standard in our analyses (see Methods), reveals associations between 13 non-covarying variants and latitude. Figure 2A shows this association for the top associated variant out of the thirteen, while Figures 2B and 2C confirm its association with latitude and flowering time. In future analyses, fine mapping of all the many combinatorial haplotypes of these 13 variants can reveal their detailed relationships to flowering time in response to photoperiod.

### References

Han, Zhongmin, Bo Zhang, Hu Zhao, Mohammed Ayaad, and Yongzhong Xing. 2016. "Genome-Wide Association Studies Reveal That Diverse Heading Date Genes Respond to Short and Long Day Lengths between Indica and Japonica Rice." *Frontiers in Plant Science* 7 (August): 1270.

Lasky, Jesse R., Emily B. Josephs, and Geoffrey P. Morris. 2022. "Genotype–Environment Associations to Reveal the Molecular Basis of Environmental Adaptation." *The Plant Cell* 35 (1): 125–38.

Yano, M., Y. Katayose, M. Ashikari, U. Yamanouchi, L. Monna, T. Fuse, T. Baba, et al. 2000. "Hd1, a Major Photoperiod Sensitivity Quantitative Trait Locus in Rice, Is Closely Related to the Arabidopsis Flowering Time Gene CONSTANS." *The Plant Cell* 12 (12): 2473–84.

Zhang, Jia, Xiangchun Zhou, Wenhao Yan, Zhanyi Zhang, Li Lu, Zhongmin Han, Hu Zhao, et al. 2015. "Combinations of the Ghd7, Ghd8 and Hd1 Genes Largely Define the Ecogeographical Adaptation and Yield Potential of Cultivated Rice." *The New Phytologist* 208 (4): 1056–66.
