## Supplementary material for "Oryza CLIMtools: A Genome-Environment Association Resource Reveals Adaptive Roles for Heterotrimeric G Proteins in the Regulation of Rice Agronomic Traits": Document S2

### Evaluation of association methods

The GWAS analysis in this study was performed using TASSEL v5.2.84 statistical software (Bradbury et al., 2007). TASSEL provides the opportunity to include indels (insertion & deletions) in the analysis, which most other GWAS software ignore. Given the presence of indels in our dataset, we used TASSEL for our study.

Before we decided on the most appropriate method to conduct GWA studies, we evaluated four models using a publicly available flowering time dataset, from which we extracted data corresponding to the landrace collection that we evaluated throughout our study: (Mansueto et al., 2017): i) a general linear model (GLM), ii) a general linear model containing a principal component analysis (PCA) correction for population structure (GLM + PCA), iii) a mixed linear model (MLM) including a kinship matrix (MLM + K) as a random effect, and iv) an MLM including both population structure and kinship relations (MLM + PCA + K). For iii and iv, the MLM used the Centered IBS matrix calculated in Tassel. GWAS results are not sensitive to the number of PCs (Price et al., 2006). For this reason, in the GLM + PCA analysis and MLM + PCA + K, we chose the first five components as the population structure matrix to conduct our GWAS (Zhao et al., 2018). The kinship matrix in the MLM models was constructed using the 'Centered IBS' kinship method in TASSEL. The MLM analysis was conducted with the options of compression and re-evaluation of variances at each marker.

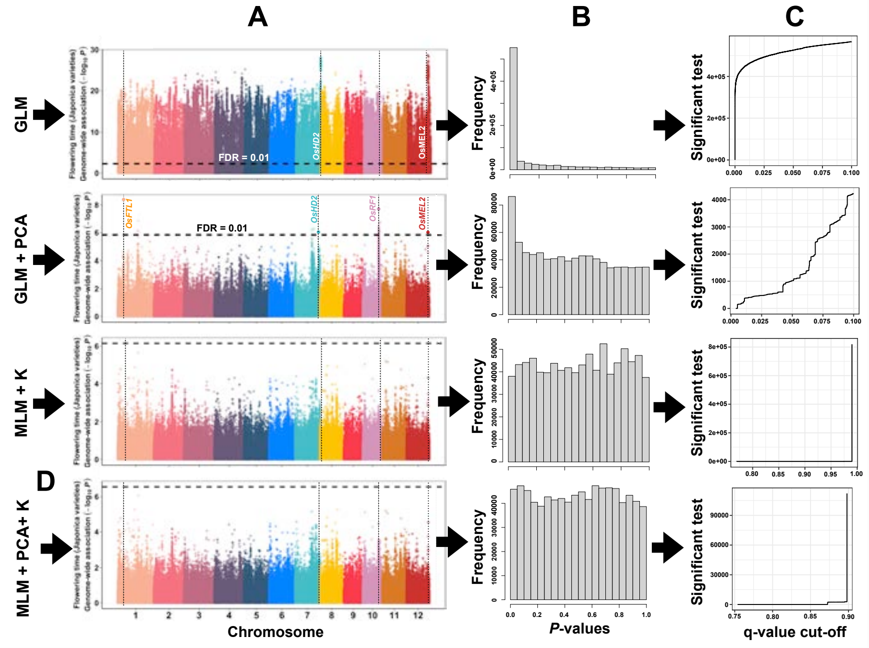


**Figure1. GLM + PCA is the most appropriate model to conduct GWAS in our dataset. A.** Manhattan plots displaying the results from four GWAS models using publicly available flowering time data. See text for a description of Manhattan plots. **B.** Distribution of *P*-values with the results of these four GWAS models. **C.** Number of significant associations obtained from these four models as a function of the q-value cut-off.

The Manhattan plots represent the *P*-values of the four GWAS models we tested (Fig. 1A). Datapoints depicting individual genetic variants are represented in genomic order by color-demarcated chromosome and position (x-axis). The value on the y-axis represents the score of the association of each variant with natural variation in flowering time for Japonica landraces. The score is the $-\log_{10}$ of the *P*-value, so the higher the score (and therefore, the lower the *P*-value), the stronger the association of any given variant with flowering time. As a result of the local correlation of causative genetic variants with neighboring variants arising from the relationship between genetic distance and recombination, groups of significant *P*-values tend to rise up on the plot, making the graph look like a Manhattan skyline, hence the name.

Flowering time, also known in rice as the heading date, is an appropriate trait to test the effectiveness of these different GWAS models in our dataset because the switch from vegetative to reproductive development is an important adaptive trait that is well-studied, and the causality of several QTLs previously identified through GWAS analysis has been demonstrated. From the study of the resulting Manhattan plots resulting from these four GWAs methods, we can identify in the GLM + PCA method four clear peaks within the FDR < 0.01 threshold in the regions of: 1) *OsFTL1* (*Os01g0218500*; *FLOWERING LOCUS T* (*FT*)-Like homolog), 2) *OsHD2* (*Os07g0695100*; *HEADING DATE 2*), 3) *OsRF1* (*Os10g0497432*; *RESTORATION OF FERTILITY 1*), and 4) *OsMEL2* (*Os12g0572800*; *MEIOSIS ARRESTED AT LEPTOTENE 2*). These genes have previously been functionally associated with the transition front vegetative to reproductive phase in rice (Komori et al., 2004; Nonomura et al., 2011; Ogiso-Tanaka et al., 2013; Liu et al., 2021), increasing confidence in the robustness of the GLM + PCA model for our application.

These peaks in the Manhattan plot are also present in the GLM model, but we cannot identify these 'towers' due to the confounding effects introduced by false positives arising from the structure of the population. Indeed, a major problem in association mapping is controlling the spurious associations that can arise from population structure. This becomes evident when we observe the distinct inflation of *P*-values that approach statistical significance in the GLM method (Fig. 1A and B). This inflation in *P*-values makes it impossible to reliably identify causal variants. The GLM with PCA model is expected to reduce false positives that arise due only to population structure (Price et al., 2006), as we demonstrate in Figure 1 for flowering time. As a result, the PCA correction for population structure allows us to identify the appropriate candidates.

The MLM with PCA and K model includes the kinship matrix in the model and is expected to further reduce false positives that arise from family relatedness (Yu et al., 2006). The MLM model alone has been reported to perform better than the GLM model alone by controlling false positives (Yu et al., 2006). The advantage of the MLM model to prevent false positives disappears for complex traits when these are associated with population structure resulting from extensive genetic divergence. The MLM model controls the *P*-value inflation well but also increases false negatives, weakening the identification of true associations (Zhang et al., 2010). Indeed, our analysis shows that the MLM model introduces an over-correction, i.e. a type II error, that renders this analysis problematic by completely removing identification of true positives. This is evident from the inspection of the Manhattan plots, where there are no significant associations using a significance threshold of FDR of 1% in the MLM models -- the four 'towers' that we identify in the GLM + PCA model, which correspond to four genes known to be flowering-related, are not present.

To further illustrate this, we applied the 'qvalue' package in R (R Core Team) to the *P*-values resulting from each of these four GWAS models to determine the number of significant associations versus each q-value cut-off (Fig. 1C). As a result, we can observe that in the GLM model, there is an inflation in the number of significant tests. This inflation is corrected once we correct for population structure (GLM + PCA). MLM models result in overcorrection and the absence of significant associations regardless of the significance threshold we impose (Figure 1C for MLM + PCA + K and MLM + K).

After careful consideration, we determined that GLM + PCA is the most appropriate model for our study. In support of this decision, previous work utilizing this GWA model in rice has proven its efficacy (Sales et al., 2017; Mogga et al., 2019; Vargas et al., 2020; Bai et al., 2022). We acknowledge that despite the imposition of correction for population structure (compared to the inflation of significant *P*-values present in the GLM model that we correct in the GLM + PCA model), our results, as is inherently true for all GWA studies, will unavoidably have a number of spurious associations. When interpreting our results, it is important to remember that correlation does not equal causation, and any associations found here cannot be assumed to be definitely adaptive or causative. We have reduced type I error, but the presence of spurious associations among the identified causative variants, while minimized, is to be expected.

MLM models are computationally demanding and require substantial computational resources and time. In our study, we conducted GWA analysis in SNPs and INDEL datasets from 658 Indica and 283 Japonica rice landraces to obtain genotype–environment associations using our dataset of 413 environmental variables. We note in passing that the computational power needed to compute the 1,656 GWAS analyses that were necessary to complete this study was alleviated by using the GLM + PCA model.
